# Bacterial GTPases act as successive placeholders to mediate ribosome assembly and its coupling to translation initiation

**DOI:** 10.64898/2026.02.15.706057

**Authors:** Aimin Cheng, Chengying Ma, Ning Gao

## Abstract

Ribosome biogenesis and mRNA translation are fundamental cellular processes regulated by a diverse set of protein factors, including GTPases. In bacteria, while several GTPases are known to participate in ribosome assembly, their precise mechanisms in subunit maturation and their potential roles in translation regulation remain largely elusive. Here, we report a series of cryo-electron microscopy (cryo-EM) structures of native pre-50S assembly intermediates and 70S translating ribosomes, isolated via epitope-tagged GTPases from engineered *Escherichia coli* cells at resolutions of 2.3-4.4 Å. These structures elucidate how three GTPases, YihA, EngA and ObgE, act as successive placeholders to mediate rRNA folding and to coordinate the correct timing of the maturation of different functional blocks within the large ribosomal subunit. Furthermore, our data identify several previously unrecognized 70S translational complexes bound by the GTPases EngA and BipA—factors traditionally regarded as assembly factors, thereby uncovering their regulatory role in bridging ribosome assembly and translation initiation. Collectively, our findings delineate a GTPase-mediated surveillance system that continuously monitors the assembly of ribosomal subunits and translation adversity, thereby safeguarding protein synthesis and maintaining proteome homeostasis.

## Introduction

Ribosomes are highly conserved, two-subunit ribonucleoprotein complex that translates messenger RNA (mRNA) into protein. In bacteria, the ribosome is composed of a small 30S subunit (SSU) and a large 50S subunit (LSU), which are responsible for decoding the mRNA and catalyzing peptidyl bond formation, respectively ^1^. Ribosome biogenesis in bacteria is facilitated by several dozens of protein factors, playing roles in the ribosomal RNA (rRNA) processing, folding and modification, and ribosomal protein incorporation ^2^. Among these biogenesis/assembly factors, GTPases consist of a particularly important group ^3,4^. Previous genetic and biochemical studies showed that depletion of individual factors often leads to growth defects, such as sensitivity to various stress conditions, and the accumulation of pre-ribosomal particles in cells ^5^. However, the understanding of their precise molecular roles in ribosome assembly has remained incomplete.

In recent years, cryo-EM studies of endogenous ribosomal assembly intermediates in eukaryotic systems, including yeast and human cells, have provided numerous high-resolution snapshots for nearly every major assembly stage ^6–10^, enabling the construction of comprehensive assembly maps for both subunits. In contrast, although *E. coli* has been a model system for ribosome assembly for over 60 years ^11–13^, our current mechanistic understanding of bacterial ribosome assembly is far from complete. For examples on the LSU assembly, previous structural studies have characterized the binding sites of several assembly factors on the mature LSU, such as GTPase EngA ^14^ and ObgE ^15^, and the low-resolution structures of assembly intermediates that accumulated upon genetic perturbations, such as depletion or deletion of specific factors (e.g., EngA, YihA, RbgA, DeaD and bL17) ^16–19^. The general assembly map for the bacterial LSU has been depicted largely from *in vitro* assembly systems ^12,20,21^. As to bacterial assembly GTPase, only ObgE has been previously employed to purify undisrupted, native pre-50S particles for structural analysis ^22^. Therefore, the absence of relevant structural information on key assembly transitions *in vivo* has precluded mechanistic understanding of how essential, energy-consuming GTPases play roles in the assembly of the 23S rRNA.

Beyond ribosome biogenesis, a few assembly factors in eukaryotes have also been demonstrated to couple ribosome assembly to translation initiation, such as eIF6 ^23,24^, Reh1 ^25^, RBFA ^26^, and MTG3 ^27^, providing an additional layer of regulation for protein synthesis. Moreover, a few translation factors, including IF3 ^28^, mtIF3 ^26^, mtIF2 ^29^, were also reported to mediate a continuum between ribosome assembly and translation. Many bacterial assembly GTPases have been linked to growth control pathways or stress responses ^3,30–33^, underlining their potential additional roles in direct regulation of translation.

In the present study, to elucidate the principles governing LSU assembly in *E. coli*, we employed a strategy using affinity-tagged assembly GTPases (YihA, EngA and ObgE) to isolate native LSU assembly intermediates for cryo-EM single-particle analysis. The resulting thirteen distinct assembly states, varying in assembly factor and ribosomal protein compositions, and rRNA conformation, provide a structural basis for the late assembly stages of the LSU. Structural analyses demonstrate that these GTPases primarily serve as placeholders that delay the pre-mature folding of a certain functional region or stabilize specific rRNA segments in distinct premature conformations, to ensure proper timing for the assembly/folding of individual rRNA domains/blocks. Moreover, we also uncover an unexpected role for EngA, probably in co-translational regulation, and establish that BipA is a *bona fide* regulatory translation factor. Together, these findings provide critical mechanistic insights into the continuous surveillance and coupling of ribosome biogenesis and protein synthesis by GTPases in bacteria.

## Results

### Cryo-EM structures of native ribosomal particles captured by epitope-tagged YihA, EngA and ObgE

To capture bacterial LSU assembly intermediates, we improved a CRISPR-based method to generate *E. coli* strains endogenously expressing FLAG-tagged GTPases (YihA, EngA, ObgE and BipA), which enable isolation of native complexes by affinity purification (**Figure S1**). Using single-particle cryo-EM, we identified twelve distinct LSU assembly states (A, B, C, D, D*, E, E*, pre-F*, F*, F, G, H) from three datasets (YihA, EngA and ObgE). Through a cascade of global and focused three-dimensional classification (**Figures S2 to S4**), these structures were determined at global resolutions ranging from 2.5 to 3.2 Å, sufficient for atomic model building of most regions (**Figure 1A**). Based on the conformational and compositional differences, these assembly intermediates could be ordered into a chronologic assembly map, manifesting that these GTPases coordinate a generally progressive conformational maturation of the 23S rRNA (**Figures 1A and 1B**). Specifically, while state A lacks the central protuberance (CP), and represents an early assembly stage similar to the “core formation” particles captured by *in vitro* reconstitution ^20^, states B-H represent late LSU assembly stages with well-folded CP, during which functionally important regions of the LSU progressively mature (**Figures 1A and 1B**). In addition to the bait GTPase, RsfS, YjgA and RluD are present in some pre-50S states, and the E-site tRNA is present in state H. Although our assignments of states A-H suggest a generally linear assembly pattern, the presence of less populated, structurally related but not identical assembly states between these described intermediates (D*, E*, pre-F*, F*) supports the notion that bacterial LSU assembly also follows parallel assembly pathways ^19^.

**Figure 1.**
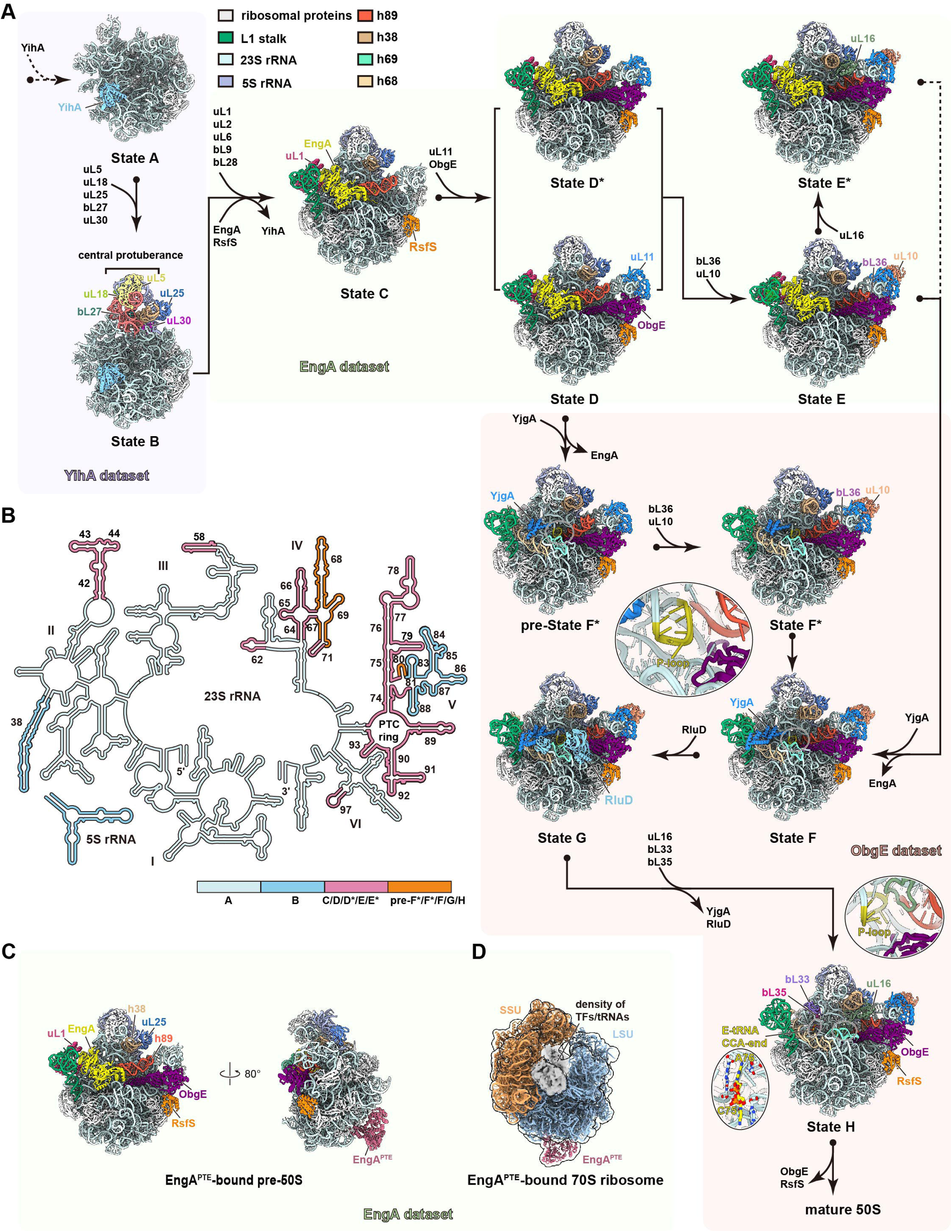
Cryo-EM structures of the endogenous ribosomal particles captured through epitope-tagged assembly GTPases. (A) Structures of the major assembly intermediates of pre-50S particles, ordered in hierarchical assembly pathways. The atomic models (States A-H) from different datasets are shown in intersubunit views, with ribosomal proteins colored in gray, and pre-rRNAs and assembly factors color-coded. Proteins that associate or dissociate between different assembly states are indicated by arrows. (B) Secondary structure diagram of the *E. coli* 23S rRNA. The rRNA elements that take turn to fold along the assembly pathways are colored-coded according to the state when they become rigid for the first time. (C) The consensus structure of pre-50S particles (EngA dataset) that also bear a second copy of EngA bound near the peptide tunnel exit (EngA^PTE^). (D) The consensus structure of the 70S ribosomes (EngA dataset) that carry EngA^PTE^. Gaussian-filtered map is superposed for clarity.

Transition from the earliest state A to the near mature state H is accompanied by sequential binding and dissociation of GTPases YihA, EngA and ObgE, concomitant with structural rearrangements of functional regions of the 23S rRNA (**Figures 1A and 1B**). While YihA binds to relatively early states A and B with large regions of the 23S rRNA in immature conformation or completely disordered, both EngA and ObgE are engaged in late assembly stages (**Figure 1B**), in line with previous studies that *in vitro* binding of EngA and ObgE could convert the mature LSU into immature conformation to certain extents ^14,15^. Importantly, the structures from the EngA dataset show simultaneous binding of EngA and ObgE in multiple states, indicating a temporal overlap of their function in coordinating the maturation of the 23S rRNA.

Very surprisingly, we also found an unexpected structural feature in two 3D classes of the EngA dataset. The first class is an immature LSU similar to state C, with EngA in its known binding position near the PTC ^14^. But different from state C, this structure also contains a second copy of EngA, occupying a position close to the peptide tunnel exit (PTE) (denoted as EngA^PTE^ for simplicity) (**Figure 1C**). The second class is the structure of translating 70S ribosomes, which also exhibits the binding of EngA^PTE^ (**Figure 1D**).

### YihA delays the folding of the L1 stalk and central helices to ensure prior installation of the CP

The CP is a key functional region of the LSU, which contributes to intersubunit interaction, tRNA positioning and movement to ensure various functional aspects of the ribosome ^34–36^. Previously, based on the structural analysis of pre-50S particles from YsxC (YihA homologue)-depleted *Bacillus subtilis* cells, it was inferred that YihA/YsxC acts to assist the maturation of the CP ^16^. However, the binding position of YihA on the pre-50S particle and its exact role remain elusive.

In our structures, a stable binding of YihA was seen in both states A and B (**Figure 2A**). While the CP is yet to be formed in state A, it has acquired its rigidity in state B with full incorporation of the 5S RNP (**Figures 2A, S5A, and S5B**), indicating that YihA accompanies the dynamic process of the CP construction. Mechanistically, YihA is docked onto a pocket exclusively formed by rRNA elements, including h33, h35a, h52 and h55/56, via a highly positively charged surface patch adjacent to its nucleotide-binding site (**Figures 2B-2D**). This binding of YihA would sterically hinder the assembly of two rRNA blocks, the L1 stalk (h75-79) and central helices (h65-71), because the assembly of them depends on their tertiary contacts with h35a and h52 from the YihA binding site (**Figures 2E and 2F**). In the mature 50S subunit, the L1 stalk base (h75, h79) needs to interact with one central helix (h66) and both of h35a and h52, and the docking of central helices also requires tertiary interactions between h65-66 and h35a/h52 (**Figures 2E and 2F**). In this way, it allows the CP block (23S rRNA domain V and 5S rRNA) sufficient degrees of freedom to fold into a stable architecture (**Figures S5A and S5B**). Given that the central helices and L1 stalk are critical structural elements for binding of translation factors and tRNAs, it could be concluded that YihA functions as a placeholder that impedes the premature formation of these two regions, ensuring prior installation of the CP (**Figure 2G**). Furthermore, ribosomal protein uL2, which contributes to formation of B7b intersubunit bridge ^37^, cannot access its docking platform due to steric hindrance imposed by YihA (**Figures 2E and 2G**). Thus, YihA also controls the timing of uL2 incorporation.

**Figure 2.**
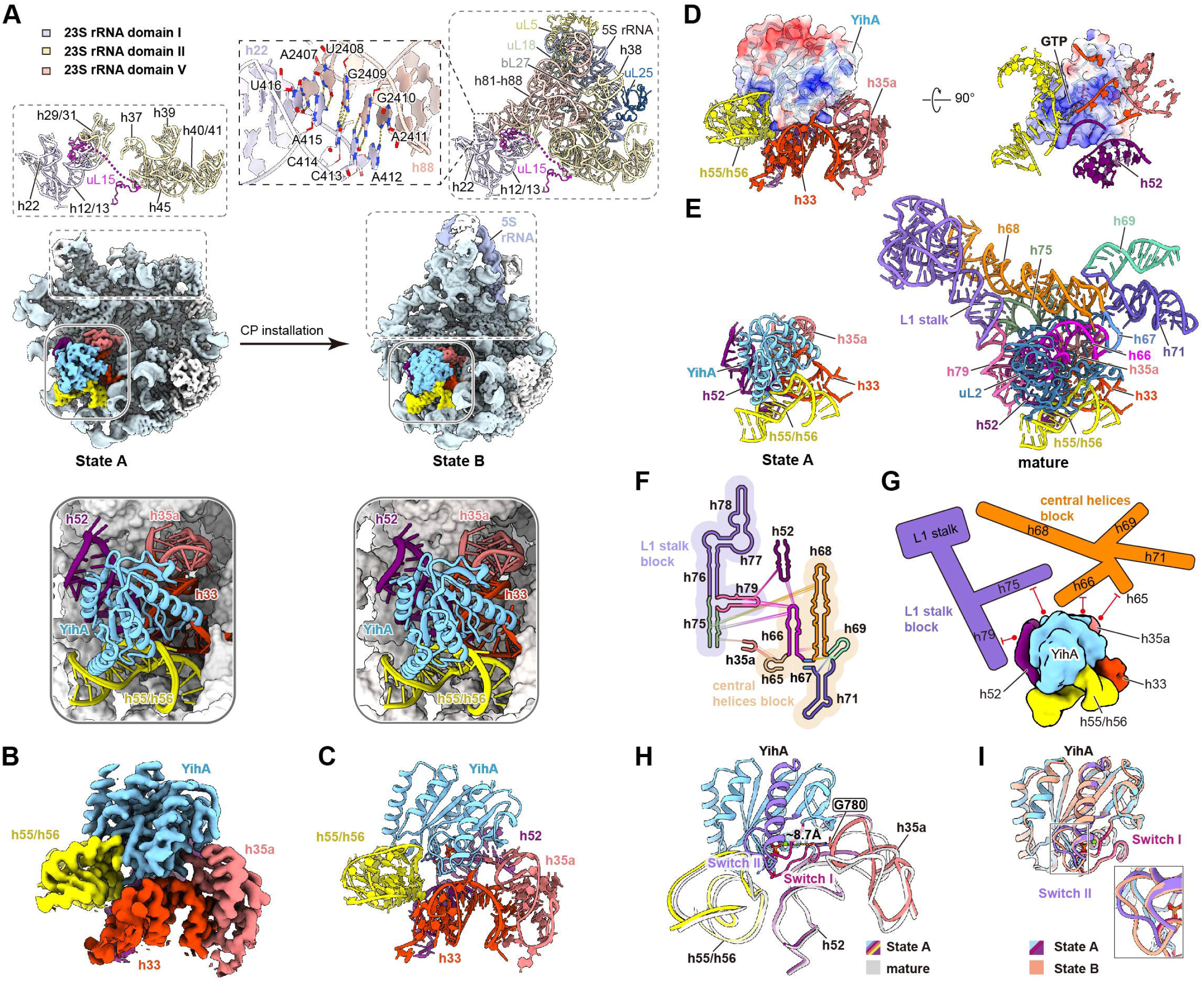
Structures of the pre-50S particles bound with YihA. (**A**) Intersubunit side views of YihA-containing States A and B (Middle panels). Top panels show the maturation of the rRNAs in the Central Protuberance (CP); bottom panels show close-up views of YihA docked on the pre-50S structures of States A and B. Proteins and rRNAs are individually color-coded. (**B and C**) Cryo-EM density (**B**) and structural model (**C**) of YihA bound to its surrounding rRNA segments. YihA, h33, h35a, h52 and h55/56 are color-coded. (D) Electrostatic surface representation of YihA at its binding site. (E) Comparison of the YihA binding site in State A with that in the mature state (PDB ID: 6PJ6). (F) Secondary structure diagram of the mature 23S rRNA, showing the tertiary contacts among the L1 stalk block (h75-79), central helices block (h65-71) and the YihA-binding rRNA helices (h35a, h52). The contacts are indicated by solid lines. (G) Schematic illustration of molecular role of YihA in preventing premature folding of the L1 stalk and central helices. (H) Conformational changes of the YihA-binding rRNA helices (h35a, h52, h55/56) between State A and the mature 50S subunit (PDB ID: 6PJ6). (I) Conformational changes in the Switch II region of YihA between States A and B.

After its initial installation, the CP undergoes a series of rotation to acquire its mature conformation ^38,39^. Comparison between states A and B shows that the nascent CP in state B is docked onto the preformed platform comprising the rRNA elements from domain I (h12/13, h22) and domain V (h29/31, h37, h39, h40/41, h45), where base pairing between h22 and h88 contributes to the stability of the CP (**Figure 2A, top**). Ribosomal proteins uL15 and bL35 were thought to be critical for initiating CP formation and positioning, respectively ^38,40^. Compared with the CP in the mature LSU, a stretch of residues of uL15 (T30-T67) remains disordered in state B and fails to provide structural support for bL35 (**Figure S5C**). Therefore, the absence of bL35 should partially account for the observed pre-mature configuration of the CP in state B (**Figure S5D**).

A general function of assembly GTPases is to act as a molecular sensor to respond to a certain assembly event to activate its GTPase activity. In the pre-50S structures, the GTPase center of YihA is exactly next to a three-way interface formed by h33/h35a, h52 and h55/56 (**Figure 2D**). To elucidate how YihA GTPase activity is coupled to a certain local conformational change during the LSU assembly, we compared the rRNA conformations of the YihA-bound state with that of the mature LSU, and identified pronounced differences in h35a and h52 (**Figure 2H**). Together with h33, these rRNA elements stabilize Switch I of YihA and hold it in a catalytically active conformation **(Figures 2H and S5E-S5G)**. Notably, G780 within immature h35a is positioned close to the GTPase center of YihA (**Figures 2H and S5H**), suggesting a potential role in activating the GTPase activity of YihA. An apparent conformational change between states A and B is in a loop of Switch II encompassing the Walker B motif (**Figures 2I and S5E**). These structural observations suggest that YihA could coordinate the correct timing of the LSU assembly events by coupling its GTPase activity with the conformational states of local rRNA elements, such as h35a/h52.

### EngA facilitates the maturation of the L1 stalk and prevents early folding of central helices h68-69

EngA (also known as Der) consists of two consecutive GTPase domains (GD1 and GD2), an RNA-binding KH domain and a C-terminal extension (CTE) ^5^. Consistent with the structures of the *in vitro* assembled 50S-EngA complex and EngA-containing pre-50S particles ^14,41^, EngA extensively interacts with the pre-50S particles (**Figure 3A**), in a position reminiscent of eukaryotic assembly factor Nmd3 ^8,42,43^. Specifically, while the GD1 anchors to the terminal region of the L1 stalk via strong interactions with uL1, L1 helices, and h88, the GD2 contacts with uL2 and attaches to h74/75 in the base region of the L1 stalk (**Figure 3B**). Notably, in the YihA-containing structure of state B, the entire L1 stalk including the base region (h74/75, h76, h79) is completely absent (**Figure 3C**). In state C, EngA, together with uL2, builds an interaction network involving bL9, h66, h58 and h21, to stabilize the three-way junction of the L1 stalk base (**Figure 3D**). This strategic position of EngA in the junction region of three rRNA blocks (the CP, central helices and L1 stalk) suggests an apparent role of EngA in facilitating the conformational maturation of the L1 stalk.

**Figure 3.**
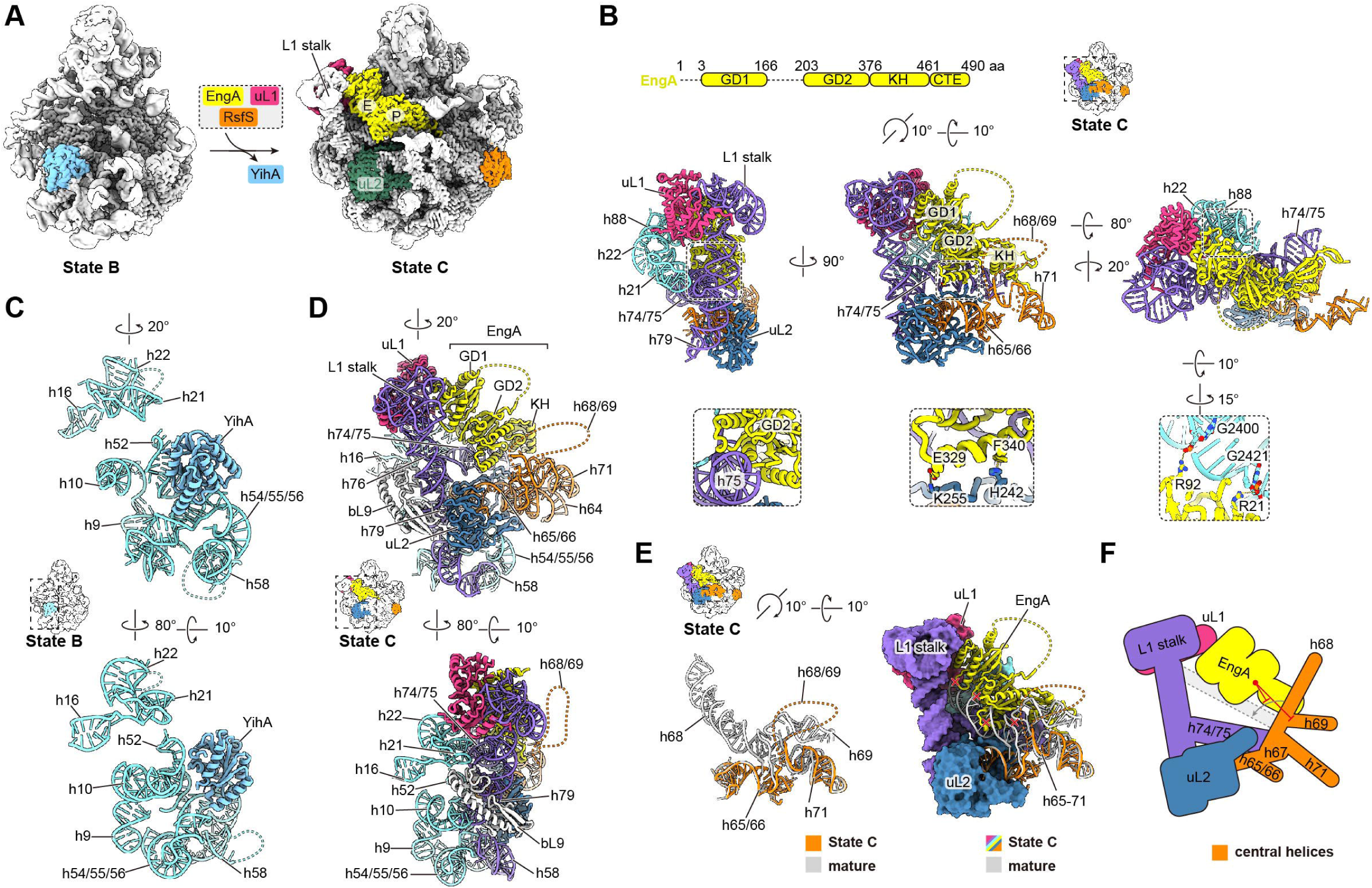
EngA coordinates the maturation of the L1 stalk and prevents early folding of central helices h68-69. (A) Cryo-EM density maps of the pre-50S structures in States B (YihA dataset) and C (EngA dataset). (B) Domain organization of EngA (top panel) and the binding site of EngA (middle panels) in State C. The specific contacts of EngA with h75, uL2 and h88, are shown in bottom zoom-in panels. (**C and D**) Comparison of the L1 stalk base region (h74/75, h76 and h79) between States B (**C**) and C (**D**), showing that the L1 stalk and helices h64-66, h71 become first ordered in State C. The rRNAs at the docking site of the L1 stalk and its base region are colored in cyan. (E) Steric clash between the central helices (h65-71) and EngA. State C is superposed with the mature 50S subunit (PDB ID: 6PJ6). Clashed sites are marked with red crosses. (**F**) Schematic model of the role of EngA in preventing the docking of the central helices h68-69.

The L1 stalk is composed of protein uL1 and h76-78 of the 23S rRNA. During translation, h78 interacts with the elbow and T/D loops of the P/E-tRNA or E-tRNA ^44,45^. Upon ribosome collision, h78 of the leading or stalled ribosome also engages with h16 of the 16S rRNA in the queueing or collided ribosome, contributing to the di-ribosome formation ^46,47^. These structural observations indicate that h78 is not merely a structural component of the L1 stalk, but a functional element of the ribosome. A recent study reported that the genes of EngA and uL1 have co-evolved with the structural changes of h78 in bacterial species ^48^, suggesting that as an assembly factor, EngA might contribute to the folding of h78. Indeed, in the EngA-containing structures of states C-E*, although the stem region of the L1 stalk, including h76-h77 is well-resolved, the h78 branch region exhibits obvious conformational heterogeneity (**Figure S6A**). Therefore, we performed focused three-dimensional classification on the region of EngA-L1 stalk using all pre-50S particles containing EngA, and resolved two slightly different conformational states for the L1 stalk, one with a disordered h78 (State 1, **Figures S6B and S6C**) and the other with a stably folded h78 (State 2, **Figures S6D and S6E**). Detailed structural analysis indicates that U2111 of h77 adopts flipped-out and flipped-in conformations in State 1 and State 2, respectively (**Figures S6C and S6E**). In State 1, U2111 interacts with a relatively conserved residue H105 of EngA probably via a water-mediated hydrogen bond (**Figures S6F and S6G**), while in State 2, U2111 is stacked against U2118 and forms noncanonical base pairing with G2144 and A2147 of h78 (**Figure S6H**), thereby stabilizing h78 in an ordered conformation. This observation suggests that the GD1 might have a direct role as a chaperone to facilitate the folding of h78.

Furthermore, EngA is embedded inside the tRNA passage encompassing the E-and P-sites, and directly interacts with h75 and h88 (**Figures 3A and 3B**). In this position, it sterically clashes with h68 in the mature 50S subunit (**Figure 3E**). As a result, the central helices h68-h69, which play important roles in intersubunit interaction and coordinating the binding and translocation of tRNAs ^49–51^, are completely disordered (**Figure 3E**). From this observation, an obvious function of EngA is to prevent the premature folding of central helices to ensure a temporal control on the assembly of different functional regions of the 23S rRNA (**Figure 3F**).

Moreover, structural comparisons of all available EngA structures in either pre-LSU-bound or free states ^52,53^ suggest that two G domains (GD1 and GD2) may function through a synergistic mechanism. In the pre-LSU-bound state, GD1 and GD2 adopt a pseudo-two-fold symmetry with Switch I in active conformation facing each other (**Figure S6I**). When in the free states, however, GD1, GD2 and KH domain display dramatically different arrangements, with Switch I in inactive conformation regardless of their nucleotide binding states (**Figure S6J**). As previously suggested ^14^, EngA activation is likely dependent on pre-50S-mediated pseudo-dimerization. During the transition from state C to state E/E*, Switch I and Switch II undergo structural rearrangements driven by the coordinated rotation of L1 stalk and the CP (**Figure S6K**). These findings suggest that the activation of GTPase activity of EngA is likely coupled to the orientation change between the CP and L1 stalk, which in turn determines the timing of EngA dissociation.

### EngA and ObgE cooperate to guide the stepwise maturation of the PTC

The peptidyl transferase center (PTC), the catalytic site of the ribosome, is constructed by rRNA elements h89-h93 of domain V and highly conserved across kingdoms of life ^54–58^. The PTC maturation is probably the most critical step, and requires the assistance of a collection of dedicated assembly factors in both prokaryotes and eukaryotes ^59–62^. Genetic data show that deletion or depletion of some assembly factors in bacteria, including those very early ones, often generate similar assembly defects in the PTC maturation ^18,19^, underlining that the PTC maturation is likely a rate-limiting step and coupled with quality control of subunit assembly. For examples, depletion of EngA or ObgE has been shown to impact the late-stage LSU assembly and delay the installation of the PTC helices ^16,22,63^. Although the *in vitro* binding properties of these two GTPases, as well as their binding sites on the 50S and pre-50S particles, have been elucidated ^14,15,22,41^, precise molecular roles and temporal sequence of their actions remain unclear.

From the EngA dataset, five distinct classes of assembly intermediates were obtained, and four of them (D, D*, E and E*) contain both EngA and ObgE (**Figures 4 and 1A**). These structures vary in protein composition of uL11, bL36, uL10 and uL16, but share common features of the immature PTC and invisible central helices. In these structures, the occupation of EngA in the tRNA passage orients its KH domain and CTE to the PTC, with the KH domain extensively interacting with h89 and h93 (**Figures 4 A-4D and S7A-S7D**). In states C and D, the binding of the KH domain to the base region of h89 generates a distinct inverted L-shape conformation for h89, compared to its mature state (**Figures 4A, 4B, and S7E)**. Since state C only contains EngA, this indicates that EngA alone is sufficient for stabilizing this pre-mature state of h89. Importantly, compared with the near mature state G, the KH domain also prevents the premature interaction between the P-loop and h89, and sterically delays the folding of the P-loop (**Figure S7F**). In the downstream state D, ObgE in the same position as previously reported ^15,22^, extends its Obg domain into the PTC from the other direction (**Figure 4B**). In this manner, the Obg domain of ObgE and the KH domain of EngA physically segregate the immature h89 from other rRNA helices, probably to prevent the stem and terminal regions of h89 from interfering with the local PTC folding.

**Figure 4.**
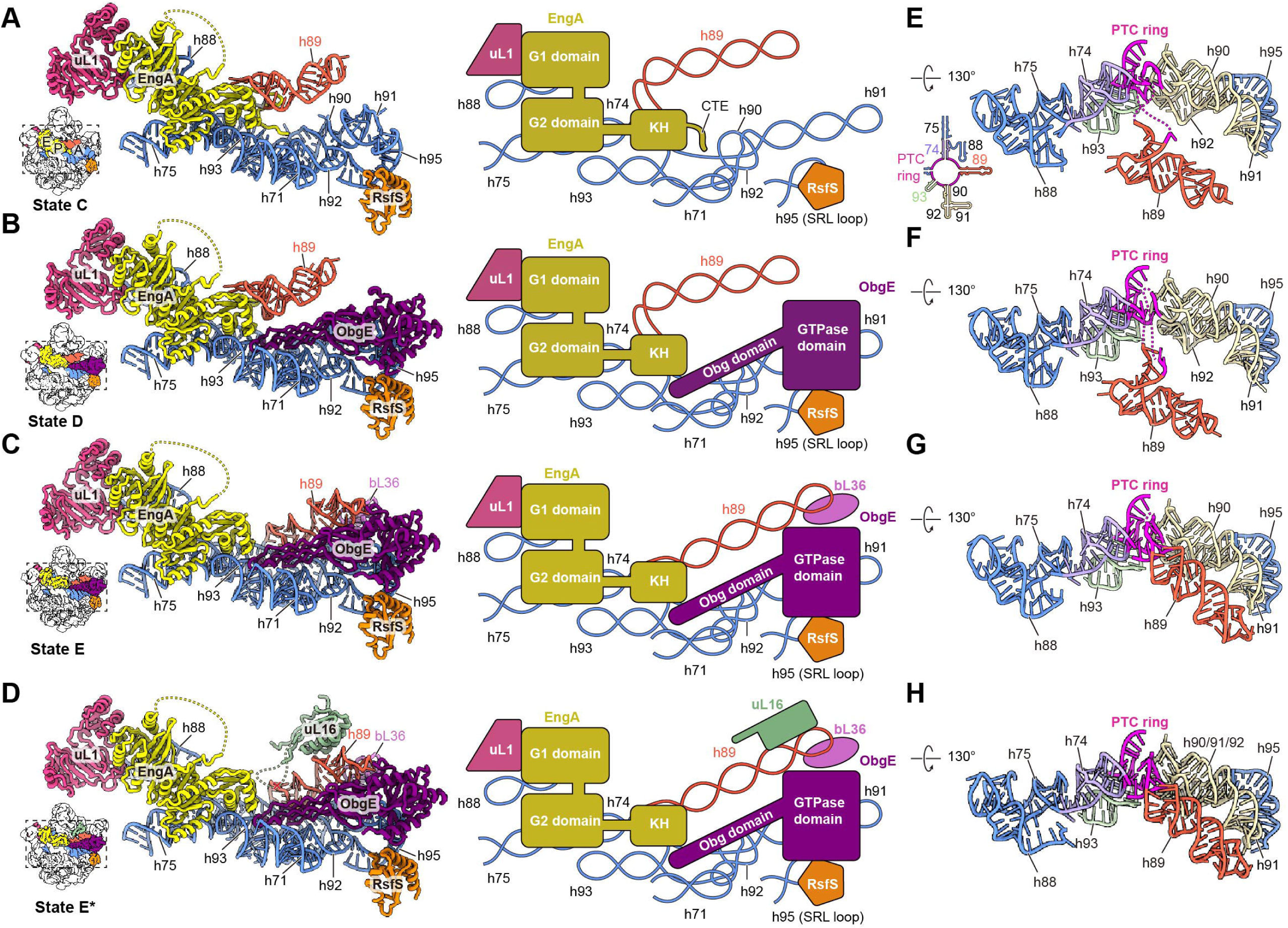
Stepwise maturation of the PTC guided by EngA and ObgE. (**A-D**) Assembly states of the PTC region bound by EngA alone or by both EngA and ObgE. State C (**A**); State D (**B**); State E (**C**); State E* (**D**). Assembly factors and rRNA helices are color-coded. Middle panels show the corresponding schematic diagrams of panels (**A-D**). (**E-H**) The conformations of the PTC helices in State C (**E**), State D (**F**), State E (**G**) and State E* (**H**) with proteins omitted for clarity.

Further structural analysis shows that simultaneous binding of EngA and ObgE facilitates stepwise conformational maturation of h89 and the PTC (**Figures 4B-4D and S7B-S7D**). From state C to state E, h89 undergoes progressive conformational changes from an inverted L-shaped structure to a near mature conformation. This structural transition of h89 is accompanied by bL36 binding in state E, and subsequent uL16 binding in state E* (**Figures 4C, 4D, S7G, and S7H**), underlining the contribution of these two proteins in stabilizing the reformed h89. Concomitantly, rRNA elements of the PTC ring (defined as G2057–C2063, G2447–C2456, C2496–C2507, G2582–G2588, A2602 and C2606–C2611 in *E. coli*) adjacent to the base region of h89, fold from disordered fragments into a well-ordered structure (**Figures 4E-4H**). These structural observations from states C to E/E* exhibit a clear pattern of stepwise conformational maturation of the PTC.

Importantly, rRNA residue-level analysis of the PTC region during these transitions reveals substantial local structural rearrangements, demonstrating that the formation of native tertiary interactions among the PTC helices (h73, h74, h89, h90/91/92, h93) involves the premature folding and subsequent unfolding of non-native structural intermediates. One such example is the junction region between the PTC and the entry of the peptide exit tunnel (PET) in the five-way junction of the PTC helices, involving G2057—C2064 from h73/h74, C2610—C2611 from h73/h93, A2503—U2504 from h89/h90, and G2446—A2448 from h74/h89 (**Figure S7I**). Among these residues, G2447, A2448, A2503, G2061 and A2062 either directly contribute to peptidyl transferase activity or are essential for polypeptide elongation ^64–68^. This PTC-PET region contains a unique helical structure initially formed in state C (**Figure S7I**). Upon the progression from state C to state G, stepwise disruption of undesired contacts and subsequent re-establishment of native tertiary interactions take place in this PTC-PET region (**Figure S7I**). Concomitant with this process, the A-site tRNA binding site, including the A-loop (U2554, C2556 in h92), together with U2492 (h89) and C2573 (h90) of the PTC helices, also acquires its native geometry (**Figure S7J**).

The results presented here indicate that the PTC region, full of tertiary interactions from multiple rRNA elements, undergoes stepwise structural transitions, and suggest that the molecular roles of EngA and ObgE are to stabilize or sequester certain pre-mature rRNA intermediates, and guide the structural maturation into coordinated and sequential steps.

### The final maturation of the central helices, the PTC and the PET are coupled

In the present study, we identified five LSU assembly states from the ObgE dataset: pre-F*, F*, F, G and H (**Figure 1A**). A previous study has also reported structural analysis of native pre-50S particles isolated through affinity-tagged ObgE ^22^. Among our structures, three states (pre-F*, F* and F) differ from previously reported states for their presence of YjgA, while state G closely resembles the reported State 3 ^22^, and state H corresponds to a near mature assembly state. Importantly, our data enable a comparative analysis of these structures with their immediate upstream and downstream assembly intermediates. The results indicate that the final maturation of the pre-50S is coordinated by a network of assembly factors, including ObgE, RsfS, YjgA and RluD (**Figure 5**).

**Figure 5.**
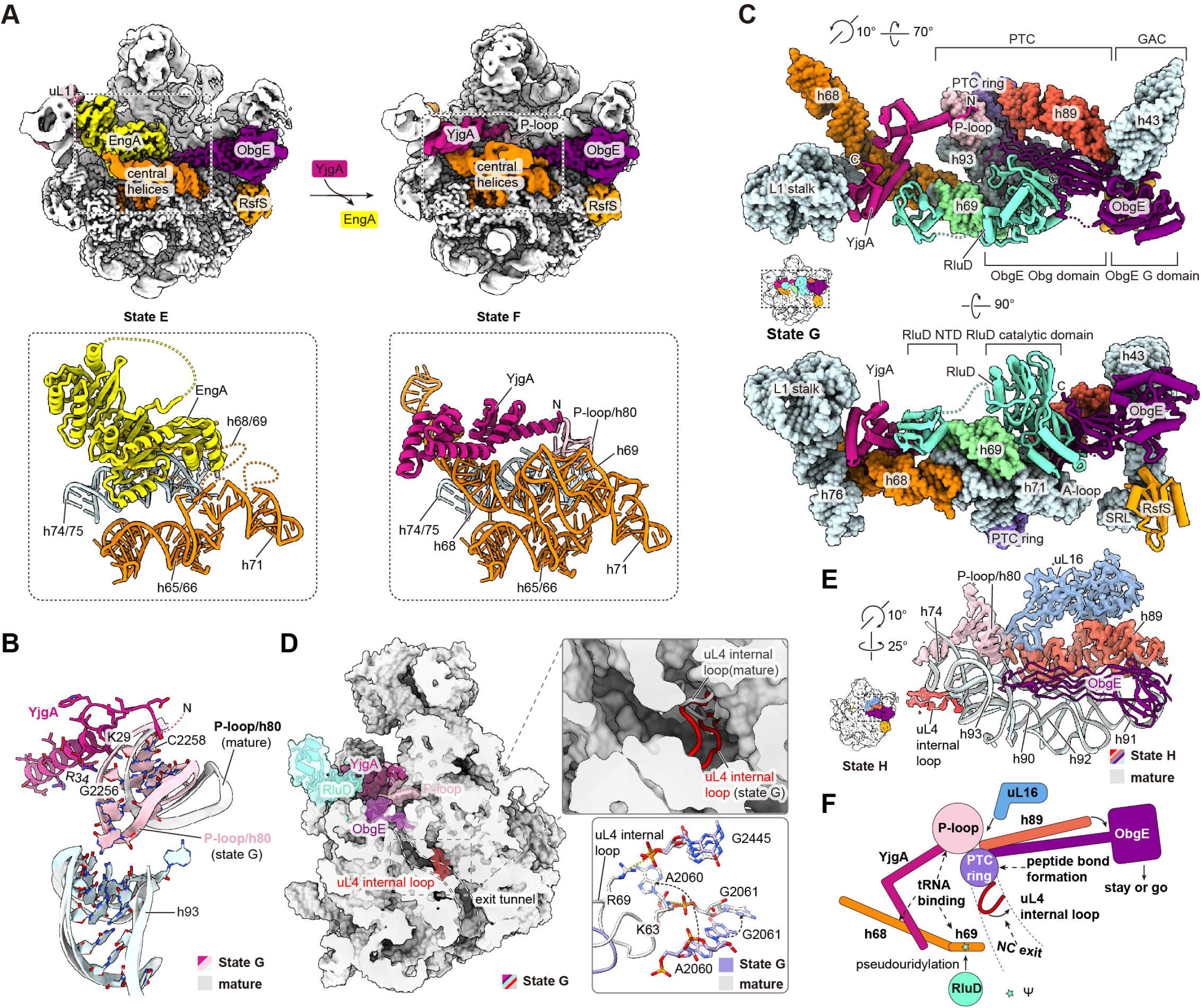
The coupled final maturation of the central helices, the PTC region and the PET. (A) Cryo-EM density maps of States E and F. EngA, YjgA and the central helices (h65-71) are shown in cartoon representation in the bottom panel. Assembly factors are color-coded individually. Central helices: dark orange; P-loop: pink. (B) The conformation of the nascent yet immature P-loop and its interaction with the N-terminus of YjgA in State G. The P-loop in mature conformation (PDB ID: 6PJ6) is superposed for comparison. (C) Interaction network of YjgA, RluD, ObgE and RsfS in State G. Assembly factors and rRNA segments are labeled and color-coded. (D) The premature internal loop of uL4 in State G. Right bottom panel shows the interaction of uL4 internal loop with the PTC-PET region in the mature 50S subunit (PDB ID: 6PJ6). (E) Comparison of the PTC-PET region in State H with the mature state (PDB ID: 6PJ6). (F) Schematic diagram of the coupled-maturation of the PTC, PET and the central helix h69 mediated by YjgA, RluD and ObgE.

In these late-stage pre-50S intermediates (pre-F*, F*, F and G), after the departure of EngA, the central helices h68 and h69 have docked onto their native positions and the P-loop has also acquired its rigidity, still in an immature conformation though (**Figures 5A and 1A**). Compared with its mature state, this nascent P-loop is oriented towards h93 (**Figure 5B**), and would sterically clash with EngA, suggesting that the formation of this initial conformation is coupled to the release of EngA. Importantly, in these structures (pre-F*, F*, F and G), YjgA docks onto the fully accommodated h68, and interacts with the nascent P-loop via its N-terminal region (**Figures 5B and 1A**). This observation indicates that YjgA only recognizes very late pre-50S particles with assembled central helices and folded yet immature P-loop.

Another functional insight of YjgA comes from state G, which contains RluD that specifically catalyzes pseudouridylation of U1911, U1915 and U1917 ^69^ within h69. RluD is embraced by two factors: while its N-terminal domain (NTD) interacts with the C-terminal domain (CTD) of YjgA, the catalytic domain simultaneously contacts both the Obg domain and CTE of ObgE (**Figure 5C**). Considering the observed conformational heterogeneity in the interface between YjgA and RluD in state G, focused 3D classification on this region was performed on all YjgA-containing particles, which revealed three kinked conformations (class 1-3) for YjgA, differing in the orientations relative to h68 (**Figures S8A-S8C**). RluD is absent in class 1 but present in classes 2 and 3, suggesting a potential role for YjgA in recruiting RluD. Electrostatic potential analyses further indicate that the orientation of YjgA is modulated by its interaction with the L1 stalk, with its CTD in class 3 showing a dramatic conformational change (**Figures S8A-S8D**). Therefore, from the observation that YjgA simultaneously interacts with the L1 stalk, h68, the P-loop and RluD, it could be inferred that YjgA might have a role in sensing the maturation status of the central helices, the L1 stalk, the PTC (h80) to ensure the timing of the modification of three functionally important residues in h69.

As to ObgE, similar to previous structures ^15,22^, its N-terminal Obg domain interacts extensively with the PTC, specifically, the PTC ring and h90/h92/h93, as well as the central helix h71, predominantly through exposed lysine or arginine residues (**Figures 5C and S8E**). The G domain docks onto h43/h44/h95, with Switch I adopting catalytically active conformation, stabilized by stacking interactions of Y189 and F191 with A1095 (h44) and A1067 (h43), respectively (**Figure S8F**). Of note, in state G that lacks uL16, bL33 and bL35, h89 exhibits a near mature conformation, and as a consequence, several key residues of the PTC-PET region (A2060, G2061, G2445) are also in immature state, which subsequently fails to support proper positioning of K63 and R69 within the uL4 internal loop, and maintains it in an immature extended conformation (**Figures 5D, S8G, and S8H**). In contrast, although state H still exhibits simultaneous binding of ObgE and RsfS, it has already assumed a mature conformation, including h89 and the uL4 internal loop (**Figures 5E and S8I**). This observation suggests that the final maturation of the PTC (h89) and PET (uL4) are tightly coupled, consistent with genetic evidence that deletion of the uL4 internal loop impairs LSU assembly ^70^.

ObgE has been proposed to function as a checkpoint factor in the final stage of the LSU assembly ^15^. Therefore, we compared states G and H to analyze the structural mechanism underlying the potential coupling between the GTPase activation of ObgE and the final LSU maturation. Comparing the conformations of h89 in states G and H suggests that transition from state G to state H would result in coordinated movement of h89, h91, h95 and β-barrel subdomain of the Obg domain, thus relieving the steric hindrance on h89 imposed by ObgE and inducing conformational changes in its G domain, particularly in Switch I and Switch II (**Movie S1**). These changes may potentially trigger its GTPase activity, ultimately resulting in the release of ObgE from the mature LSU. This GTPase-driven placeholder behavior of ObgE in state G—holding near mature H89 on the immature LSU and in state H—completing H89 and PTC maturation, closely resemble the sequential actions of GTPBP10 ^71^ and GTPBP5 ^72^, two ObgE homologs, in mitochondrial LSU assembly (**Figure S8J)**.

Collectively, the structural findings from the ObgE dataset reveal that the final maturation steps of the functional regions, including the central helices, PTC and PET are tightly coupled by an interaction network of assembly factors (**Figure 5F**). Particularly, YjgA couples the maturation of the distal central helix h68 to the PTC region, whereas ObgE licenses the final properly assembled LSU for entry into the translating pool.

### A second role of EngA in co-translational regulation

The presence of EngA^PTE^ in pre-50S and 70S structures from the EngA dataset (**Figures 1C and 1D**) prompts us to further characterize the structural features of these two populations. First, based on the occupancy of EngA^PTE^, we classified all the pre-50S particles from the EngA dataset into two groups (**Figure S3C**). The consensus maps of them indicates that besides the presence or absence of EngA^PTE^, the two structures are nearly identical, with similar pre-mature features on the pre-50S particles (**Figure 6A**), suggesting that this second binding site of EngA is independent of its known role in the PTC maturation. Second, through 3D classification of all the 70S particles from the EngA dataset, we isolated a subset of 70S particles with the binding of EngA^PTE^ (**Figures 6B and S3C**). The consensus map of these 70S particles contains unassigned densities in the intersubunit space, presumably corresponding to certain translation factors and tRNAs. This consensus map also has mRNA density in the mRNA channel of the SSU, and nascent chain density in the PET of the LSU (**Figures 6C and 6D**), indicating that these 70S particles are engaged in active translation. Further focused 3D classification on the intersubunit space revealed three major populations for these 70S ribosomes, with each of them bound with a different factor, EF-G, EF-Tu or BipA, a functionally enigmatic GTPase (**Figures 6E-6G**). These results indicate that EngA^PTE^ is capable of associating with translating ribosomes, and highlight a potential role of EngA^PTE^ in co-translational regulation.

**Figure 6.**
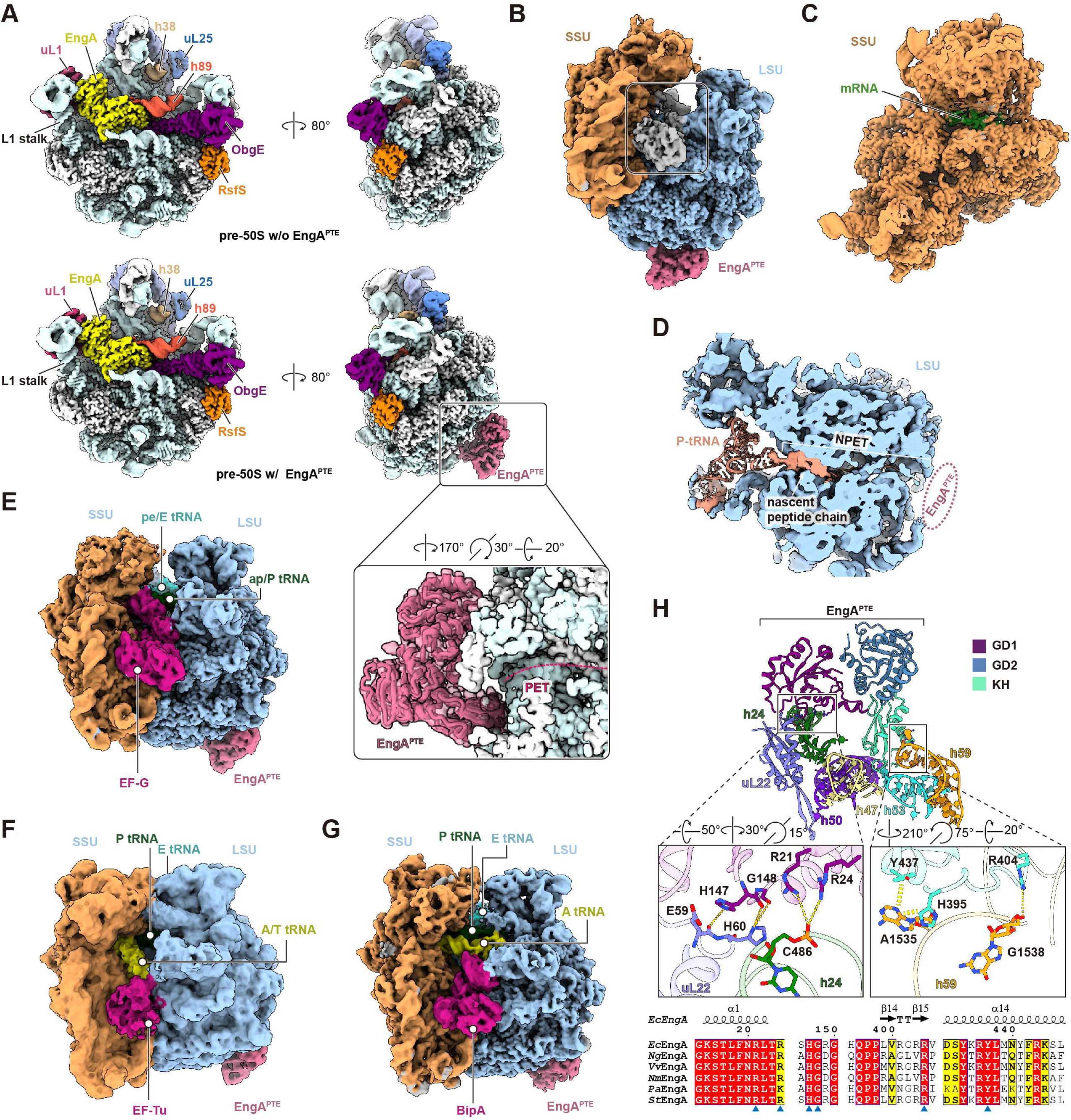
Structures of the pre-50S and translating ribosomes with the binding of EngA near the peptide tunnel exit. (A) The consensus cryo-EM maps of the pre-50S particles with or without the binding of EngA^PTE^. Factors and rRNA are individually color-coded. (**B-D**) The consensus cryo-EM map of the EngA^PTE^-containing 70S ribosomes (**B**), with the mRNA (**C**) and nascent peptide chain (**D**) densities highlighted. (**E-G**) Cryo-EM reconstructions of the EngA^PTE^-containing 70S ribosomes with EF-G (**E**), EF-Tu (**F**) and BipA (**G**) bound. (**H**) Interactions of EngA^PTE^ with the pre-50S particles. Close-up views of the interface are depicted in the middle panels. Bottom panel shows the sequence alignment of the EngA^PTE^ interface residues. *Ec*, *E. coli*; *Ng*, *N. gonorrhoeae*; *Vv*, *V. vulnificus*; *Nm*, *N. meningitidis*; *Pa*, *P. aeruginosa*; *St*, *S. Typhimurium*. Conserved residues are indicated by blue solid triangles.

Interestingly, EngA^PTE^ adopts a conformation distinct from that of EngA^PTC^, characterized by an approximate 30° rotation of the GD1 domain relative to the GD2 and KH domains (**Figures S9A and S9B**). Notably, the Switch I regions of both GD1 and GD2 in EngA^PTE^ also exhibit dramatic conformational differences, and appear to be in catalytically inactive state (**Figure S9C**). EngA^PTE^ specifically interacts with ribosomal components through its GD1 and KH domain (**Figure 6H**). The GD1 is docked onto h24 and uL22. Basic residues R21 and R24 of EngA form electrostatic interactions with the phosphate group of C486 in h24; H147 and G148 of EngA contact E59 and H60 of uL22 via hydrogen bonds. The KH domain interacts with h59, with R404 contacting the phosphate group of G1538 and H395/Y437 forming hydrophobic interactions with the base of A1535. Most of the interface residues of EngA are highly conserved across Gram-negative bacteria (**Figure 6H**), suggesting a conserved and potentially universal role for EngA^PTE^ at the PTE.

It is well known that h59 plays an important role in mediating the interactions of various co-translational factors, including SecYEG, YidC, Signal Recognition Particle (SRP), Trigger Factor (TF) and SecA, to facilitate co-translational nascent polypeptide processing, folding and translocation ^73–76^. Similar to these h59-interacting factors ^77,78^, the binding of EngA^PTE^ also induces a tilted conformation for h59 (**Figure S9D**). Importantly, EngA^PTE^ sterically clashes with all these co-translational factors at the PTE (**Figures S9E and S9F**). Given the observation that EngA^PTE^ binds to both late-stage LSU intermediates and 70S ribosomes, it is highly likely that EngA^PTE^ mediates a continuum between LSU assembly and translation initiation. This is consistent with the emerging concept that ribosome assembly and translation initiation are tightly coupled, and the final release of several assembly factors has been demonstrated to take place during the initiation and early elongation phases ^24–27,29,79^.

### BipA is a *bona fide* regulatory translation factor

Different from EngA, BipA is an elongation factor-like GTPase and contains a characteristic histidine residue at its catalytic site for GTP hydrolysis (**Figure S10A**), a defining hallmark of translational GTPases. Previously, a medium-resolution (4.7 Å) cryo-EM structure of an *in vitro* assembled 70S-BipA complex has been reported, showing that BipA could engage with the ribosome bearing both A- and P-site tRNAs^80^. However, functional data showed that deletion of *bipA* results in a cold-sensitive phenotype associated with LSU assembly defects, and this phenotype could be rescued by the loss of RluC (a pseudouridine synthase that modifies U955, U2504, and U2580 of the 23S rRNA) or exacerbated by the further deletion of *deaD* (a DEAD-box RNA helicase that facilitates the folding of the 23S rRNA), suggesting a role of BipA in the LSU assembly ^81,82^. Therefore, the precise physiological role of BipA has remained contentious, with the debate centering on whether it functions as an assembly factor or a translation factor ^81,83–85^.

The simultaneous presence of BipA and EngA on the 70S ribosome (**Figure 6G**) suggests that they may function together during the early stages of translation. This unexpected finding prompted us to characterize the structures of endogenous BipA-bound ribosomal particles. Towards this purpose, a knock-in *E. coli* strain with BipA epitope-tagged at the N-terminus was constructed (**Figure S10B**). Sucrose gradient fractionation and pull-down assays demonstrated that BipA predominantly associates with 70S ribosomes and polysomes (**Figures S10C and S10D**). Consistently, 3D classification of the BipA dataset revealed that approximately 94% of 70S ribosomes were bound with BipA (**Figures 7A and S11**). The final density map of the BipA-bound ribosome displays well-resolved density for all translational components, including three tRNAs in the A-, P- and E-site, and a fragment of the mRNA (**Figures 7B and 7C**). Moreover, the nascent chain density in the PET is also clearly tracible (**Figure 7D**). These observations indicate that BipA is engaged with translating ribosomes.

**Figure 7.**
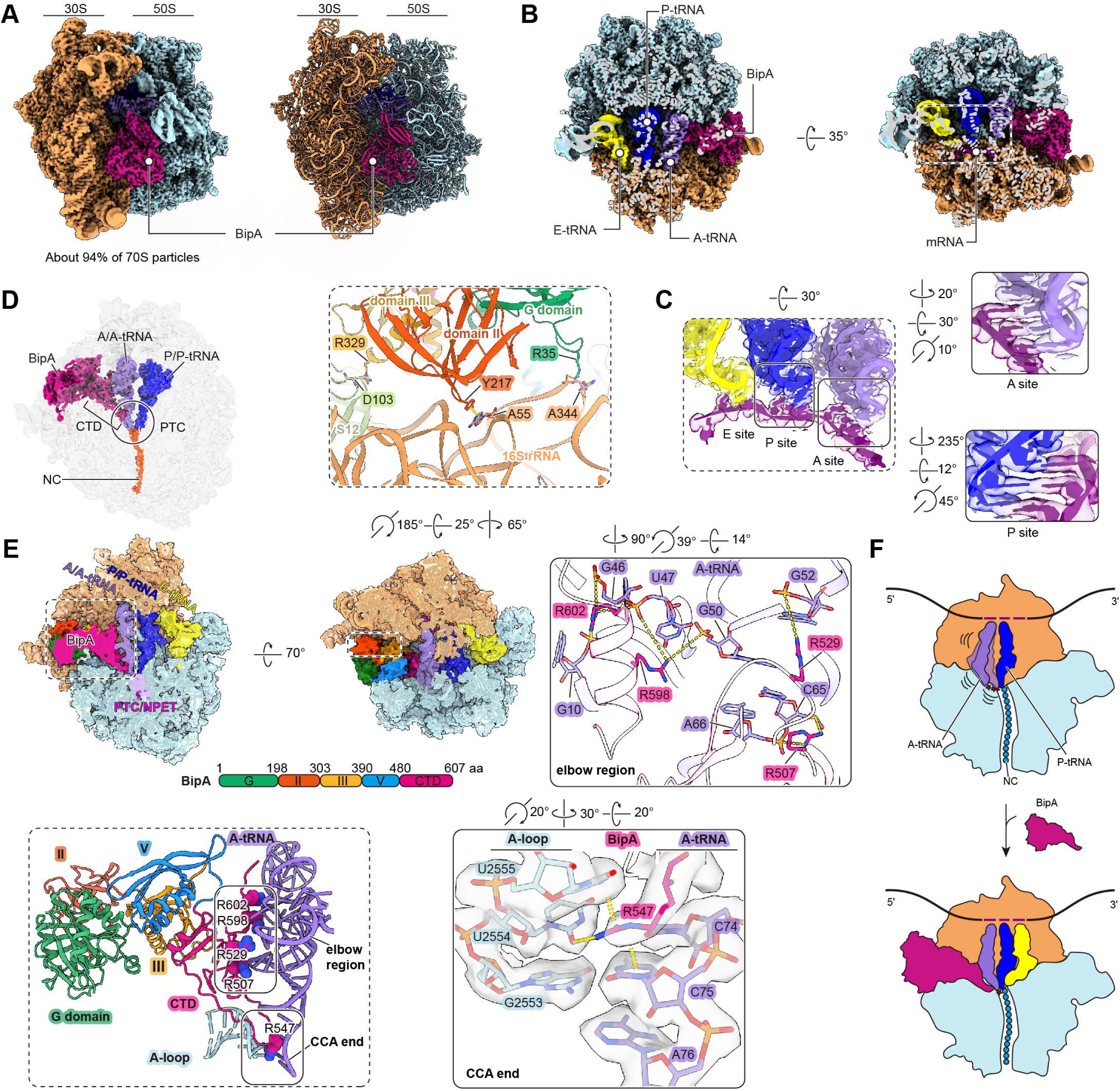
Structure of the BipA-bound translating ribosome. (A) The cryo-EM map and atomic model of the 70S-BipA complex. (B) Cross-section view of the cryo-EM map, showing the presence of mRNA and three tRNAs in the 70S-BipA complex. (C) Zoom-in views showing the tRNA-mRNA duplexes in the A and P sites. (D) Segmented cryo-EM densities of the A-tRNA, P-tRNA, nascent peptide chain (NC) and BipA are highlighted. (E) Interaction of BipA with the A-tRNA and the 30S subunit. Interaction interfaces are shown in three zoom-in views. (F) Schematic diagram illustrating BipA-mediated stabilization of the A-tRNA during translation.

The A- and P-site tRNAs adopt the standard A/A and P/P configurations, and exhibit canonical base-pairing interactions with the mRNA (**Figures 7C and 7D**). Because the proper orientation of the 3’-CCA end of the A-tRNA is crucial for peptide bond formation, we analyzed whether BipA stabilizes the 3’-CCA end of the A-tRNA in a proper geometry. BipA shares structural similarity with EF-G and LepA ^80^, but features a unique C-terminal domain (CTD) that is positioned near the ribosomal A site to extensively interact with the A-tRNA (**Figure 7E**). Compared to the canonical pre-translocation state ^58^, the A-tRNA is distorted with marked displacements in the elbow region and the acceptor arm when the anticodon stem-loop is used as reference of alignment (**Figures S12A and S12B**). Despite these large distortions, however, on the 70S ribosome, the CCA-end of the A-tRNA is accessible to that of the P-tRNA, with a comparable distance to the CCA-end of the P-site tRNA as seen in canonical pre-translocation complex (**Figure S12C**). This configuration contrasts with the dramatically disrupted geometry of the A-tRNA induced by LepA ^86^, another paralog of EF-G (**Figure S12C**). The high-resolution cryo-EM map of the BipA-bound ribosome enables unambiguous modeling of the unique CTD of BipA. Specifically, the CTD interacts with elbow of the A-tRNA via several conserved arginine residues (R507, R529, R598 and R602) (**Figures 7E and S12D**). And very interestingly, R547 of BipA, which is located at the tip of a long loop (residues 536-564) in the CTD, occupies the position of C74 of the A-tRNA, resulting in a rotation of the C74 base (**Figures 7E and S12E**). Although the noncanonical base pair between C74 of the A-tRNA and U2554 of the A-loop is disrupted in this configuration, R547 appears to potentially compensate for the disruption by forming stacking interactions with C75 of the A-tRNA and U2555 of the A-loop, as well as hydrogen-bond interactions with U2554, thereby maintaining 3’-CCA end of the A-tRNA in a translation-favored geometry (**Figures 7E and S12E**).

To assay the functional implications of the interactions between BipA and ribosomal components, we introduced BipA mutations into different interfaces. First, at the interface of the elbow region and CCA-end of the A-tRNA, single point mutation of R547A or quadruple mutations of R507A, R529A, R598A and R602A both resulted in growth defect at low temperature (**Figures 7E and S12F**). Second, a triple mutation (R35A, Y217A, R329A) that disrupts the interactions between BipA and the 30S subunit also caused slow growth under cold stress (**Figures 7E and S12F**). As a control, similar to previous work ^87^, growth defect was also observed for GTPase-dead mutant (H78A or H78E) (**Figure S12F**). These results confirm that the function of BipA is dependent on its interactions with translating ribosomes and mechanistically coupled to GTP hydrolysis.

Since the CCA-end of the A-tRNA in the BipA-bound ribosome appears to be in a conformation compatible with peptide bond formation, we thus investigated the interplay between BipA and other non-canonical translation factors involved in resolving poly-proline-dependent stalling, including UUP ^88^, EF-P ^89,90^, and YeiP ^91^. Spot assays revealed that only UUP, a member of the ATP-binding cassette family F (ABCF), could rescue the cold-sensitive growth defect of Δ*bipA* strain (**Figure S12G**). UUP features a P-site tRNA Interaction Motif (PtIM) that stabilizes the acceptor stem of the P-tRNA in a favorable geometry for peptide bond formation, highly similar to that observed in the presence of EF-P, to resolve ribosome stalling ^89,92–94^. We next introduced mutations to a few PtIM residues (R271A, K275A, R277A and R278A) that directly interact with the P-tRNA and the 23S rRNA. This mutant was unable to complement the defect of Δ*bipA* strain (**Figure S12H**). Therefore, these results suggest that BipA might have a partial functional redundancy with UUP to alleviate certain translational stalling situations. The resolution of this type of stalling could be achieved either by UUP-mediated P-tRNA stabilization or by BipA-mediated A-site tRNA stabilization (**Figure 7F**).

## Discussion

Ribosome biogenesis and translation are tightly orchestrated by a collection of ribosome-associated GTPases. In this study, we characterized the structures of endogenous ribosomal particles affinity-purified using some of these GTPases as baits, including YihA, EngA, ObgE and BipA. These high-resolution snapshots provide structural and mechanistic insights into GTPase-mediated maturation of the LSU and translation regulation in bacteria.

First, our structural analyses show that assembly GTPases, such as YihA, EngA and ObgE, play important roles in orchestrating key assembly events of the LSU, acting as placeholders to ensure an ordered conformational maturation of different functional regions, including the CP, L1 stalk, central helices, and the PTC (**Figure 8**). Specifically, YihA, as the earliest binding GTPase among the three, blocks the pre-mature folding of the L1 stalk and central helices to ensure the prior installation of the CP. Subsequent dissociation of YihA is accompanied by the binding of EngA, which acts in turn as a placeholder to prevent the pre-mature docking of central helices, but allows the CP to undergo further maturation steps. After the binding of ObgE, EngA and ObgE together orchestrate the stepwise maturation of the PTC (including the PTC-ring, A-loop and h89). The departure of EngA enables the docking of h68 and other central helices, which provide a binding platform for YjgA and RluD, allowing continuous subtle maturation of the PTC region. The final maturation of the PTC-PET junction (including the constriction loop of uL4) is monitored by ObgE and coupled with its release. As illustrated in this assembly map (**Figure 8**), a principal function of these GTPases is to establish the correct temporal sequence for the assembly of different regions.

**Figure 8.**
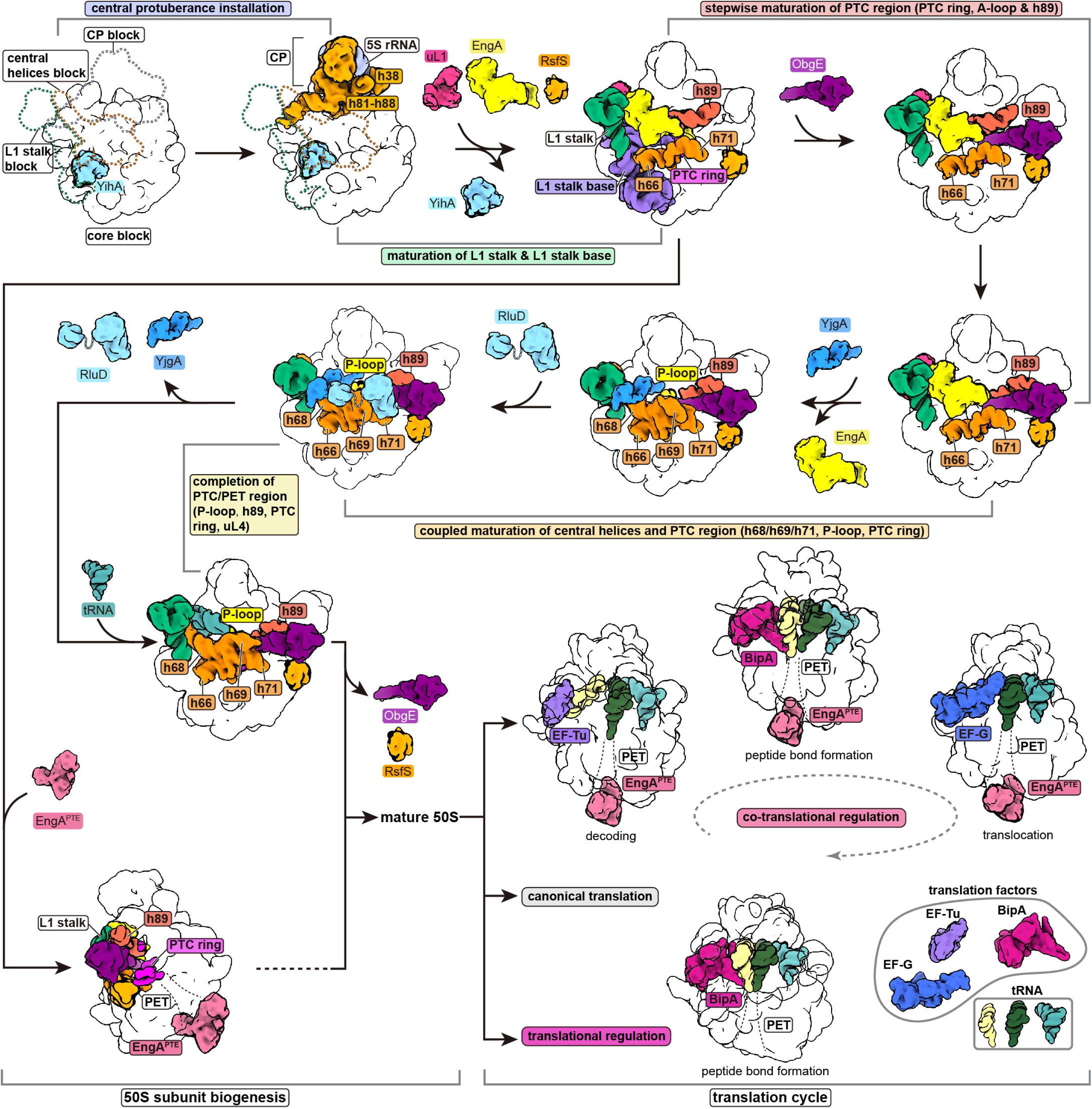
Model for the GTPase-mediated temporal maturation of the LSU and the continuum between ribosome assembly and translation. Simplified schematic model illustrating GTPases (YihA, EngA and ObgE) as placeholders to mediate temporal organization of functional regions of LSU, and GTPases (EngA and BipA)-mediated continuum between ribosomal assembly and translation in bacteria. Key transitions, including the installation and maturation of functional centers (the CP, L1 stalk, central helices, PTC ring, h89, A/P-loop and PET), as well as GTPases (EngA and BipA)-mediated translation regulation, are indicated.

Second, our data also show that EngA might have a previously unrecognized role in translation regulation. EngA^PTE^ is in conflict with many co-translational factors binding at the PTE of the ribosome, including SecYEG translocon, SRP, TF and SecA, indicating that a sustained association of EngA^PTE^ at the PTE might interfere with co-translational events. We propose that EngA^PTE^ may function through several possible mechanisms: (1) The absence or presence of EngA^PTE^ is employed to regulate the subcellular distribution of 50S subunits and 70S ribosomes. EngA^PTE^ would prevent the attachment of the ribosome/50S subunit to the cell membrane and maintain its cytosolic localization. Very recently, Lsg1, an LSU assembly GTPase in eukaryotes, has been proposed to function in a similar manner to regulate the endoplasmic reticulum (ER) localization of 60S and 80S ribosomes to modulate translation ^95^. (2) EngA^PTE^ regulates co-translational translocation of certain outer membrane proteins (OMPs). The incompatibility of EngA^PTE^ with TF at the PTE suggests that EngA^PTE^ could modulate the accessibility of nascent chains, and may function as a TF surrogate. TF functions as a major co-translational chaperone, and together with SRP, accurately sorts nascent proteins into both co-translational and post-translational targeting pathways ^96,97^. In particular, TF is enriched on ribosomes translating OMPs ^98^. Notably, EngA is co-transcribed with BamB in the same operon, which encodes a subunit of the β-barrel assembly machinery (BAM) complex required for the biogenesis of OMPs in Gram-negative bacteria. Therefore, this genetic arrangement raises the possibility that EngA, analogous to TF, may contribute to the translocation of OMPs. (3) EngA^PTE^ inhibits various co-translational processes to downregulate translation in response to stress signals. It is known that cellular concentrations of different nucleotides, as a function of growth phase and metabolic state, play important roles in bacterial physiology. The fact that *E. coli* growth rate correlates well with intracellular EngA abundance ^32^, suggests that EngA might be one of candidates to sense the cellular GTP/GDP ratio. The overall conformation of EngA ^99^, as well as its binding to different ribosomal fractions ^100^, could be fine-tuned by nucleotide-binding states of its two GTPase domains. Since the two GTPase domains of EngA differ in nucleotide affinity ^53,100,101^ and GTP-hydrolyzing activity ^53,102^, this would make EngA an ideal effector protein that could sense a wide-range change of the GTP/GDP ratio in the cell. In addition, EngA also binds to the alarmone ppGpp *in vitro* ^103^ and has been functionally linked to the stringent response pathway in *E. coli* ^31^. It was also shown that overexpression of RelA, the enzyme responsible for (p)ppGpp synthesis, rescues the growth defect of an EngA (N321D) mutant strain ^31^. Therefore, these data suggest that EngA^PTE^ might act to downregulate translation, particularly the co-translational events, in response to growth control signal.

Third, the temporal overlap of EngA^PTE^ and BipA in translating ribosomes supports a *bona fide* role of BipA in translation regulation. BipA may function as a component of a translation surveillance system that includes other noncanonical translation factors, such as EF-P and UUP, to resolve different stalling events, as deletion of *bipA* exacerbates the cold-sensitive effect in the Δ*efp* strain ^93^ and overexpression of UUP rescues the defects of the Δ*bipA* strain (**Figures S12G and S12H**). Given that translation initiation and early elongation phase are rate-limiting steps for protein synthesis ^104,105^, BipA may function in early phases of elongation. In particular, BipA and UUP may function in a synergetic manner to stabilize the CCA-ends of the A- and P-site tRNAs, respectively, to improve the efficiency of certain events of peptide bond formation. Another supporting evidence is that disruption of pseudouridine synthase RluC also suppresses the defects of the Δ*bipA* strain ^81^. The nucleotide substrates of RluC, U955, U2504, and U2580 of the 23S rRNA, are exactly located close to the interface between the BipA CTD and the A-site tRNA. And an additional piece of evidence comes from the observed densities of the Shine-Dalgarno (SD) and anti-Shine-Dalgarno (aSD) duplex and bS21 in the 70S-BipA complex (**Movie S2**), strongly suggesting the engagement of BipA in translation initiation or early elongation.

In summary, our study provides a structural basis for understanding the molecular roles of several assembly GTPases in the maturation of LSU in bacteria, and more importantly, uncovers unexpected roles of EngA and BipA in translation regulation. These data show that late-stage assembly factors, such as EngA, mediate a continuum between ribosome assembly and translation initiation. As the ribosome is one of the most important targets of antimicrobial agents, with many clinically used antibiotics engaging conserved sites within the PTC and the PET ^106^. The knowledge we learned from these GTPase-mediated regulatory events might be harnessed for rational design and development of next-generation antimicrobial compounds.

## Data availability

Cryo-EM density maps have been deposited in the Electron Microscopy Data Bank (EMDB) under accession codes EMD-xxxxx (). The corresponding atomic models have been deposited in the RCSB Protein Data Bank (PDB) under accession codes xxxx ().

## Acknowledgements

We thank Prof. Jianguo Yang (Peking University) for sharing plasmids for lambda red recombination system; Prof. Jian Lu (Peking University) and Xinyue Chen, Peixiang Gao for help with biochemical assay of BipA; Yang Liu for help with reagents; the Core Facilities at School of Life Sciences, Peking University for assistance with negative staining electron microscopy; the National Center for Protein Sciences at Peking University for assistance with imaging facilities; the Cryo-EM Platform and the Electron Microscopy Laboratory of Peking University for help with cryo-EM data collection; the High-Performance Computing Platform of Peking University for support with computation. This work was supported by National Key R&D Program of China (2024YFA1306202 to N.G.) and the National Science Foundation of China (32230051 to N.G.).

## Author contributions

N.G. conceived and supervised the project. A.C. performed molecular cloning and genome editing, purified ribosomal samples, and prepared grids for cryo-EM study. A.C. conducted cryo-EM data collection and image processing. C.M. calculated cryo-EM maps of states A and B from the YihA dataset. A.C. built and refined the atomic model with assistance from N.G. A.C. performed the spotting assays. A.C. and N.G. analyzed the data. A.C. prepared the figures and drafted the manuscript. N.G. and A.C. revised the manuscript.

## Materials and Methods

### Bacterial strains and culture conditions

*E. coli* strains DH5α and BW25113 were used for plasmid construction and ribosome purification, respectively. All bacterial strains were grown in Luria-Bertani (LB) media or on Agar-solidified LB media. Kanamycin (30 μg/mL), Ampicillin (100 μg/mL), Streptomycin (50 μg/mL), Chloramphenicol (34 μg/mL) and L-arabinose (25 mM) were added to culture media when required.

### Plasmid construction for genome editing

The pKD46-FnCpf1 and pAC-crRNA vectors were constructed as previously described ^107^. For donor DNA, homologous regions (500-800 nucleotides) flanking the genomic insertion site, and the in-frame coding sequence of StrepII-TEV-3xFLAG or 3xFLAG-TEV-StrepII fused with a kanamycin resistance cassette (FRT-kan^R^-FRT) derived from plasmid pKD4 ^108^, were cloned into pUC57 (GenScript) to serve as the template for preparation of linear donor DNA for genomic insertion. For guide RNA (CrRNA), each sequence targeting the gene of interest was designed using the online tool (https://crispor.gi.ucsc.edu) and subsequently cloned into the pAC-crRNA vector according to previously established protocols ^109,110^.

### Generation of knockin and knockout strains

Genome-edited strains were generated as previously described with modifications ^107^. Briefly, the insertion sequences and flanking homologous regions were amplified using PrimeSTAR Max DNA Polymerase (Takara). The resulting PCR products were purified and denatured through a “fast cooling” procedure. Denatured linear donor DNAs and their respective guide RNA plasmids were co-introduced into electrocompetent BW25113 cells ^108^ to facilitate genomic integration. Following electroporation, cells were recovered for 2-5 hours in the presence of 25 mM L-arabinose and plated on LB agar medium supplemented with 30 μg/mL kanamycin, 34 μg/mL chloramphenicol and 50 μg/mL streptomycin. After 2 days of incubation at 30□, positive clones were screened by PCR and verified by Sanger sequencing. The strains of interest were then transformed with pCP20 ^111^, which encodes FLP recombinase (FLPD5 variant) from *Saccharomyces cerevisiae* to excise the kanamycin resistance gene flanked by FRT sites. Strains were cured of pCP20 by incubation at 42□, and successful genomic integrations were confirmed by PCR and Sanger sequencing. Gene knockout was performed following the same procedure, with open reading frame replaced by kanamycin resistance cassette in the donor DNA instead.

### Purification of native ribosomal particles

Eight liters of LB cultures as a batch unit were inoculated with genomic GTPase-tagged strains from a stationary culture to a starting OD600 of ∼0.05. Cells were cultured at 37□ unless otherwise specified and agitated at 220 rpm to an OD600 of 0.8-0.85. Cells were harvested using a JLA8.1000 rotor (Beckman Coulter) for 10 minutes at 5,000 rpm and 4□, then washed once in fresh LB medium. Pellets were flash-frozen in liquid nitrogen and stored at -80□. The cell pellets were resuspended in lysis buffer (50 mM HEPES pH 7.5, 75 mM NH_4_Cl, 20 mM MgCl_2_, 25 μL RNase-free rDNase I, 25 μL RiboLock RNase inhibitor, 5 μg/mL lysozyme, 0.9 mM GDP•BeF_3_^-^, 1x protease inhibitor cocktail, 1 mM PMSF) on ice and lysed by low-power (30%) sonication. The lysate was centrifuged in Eppendorf Centrifuge 5804R for 15 minutes at 10,000 rpm and 4□. The resulting supernatant was transferred to a fresh tube and subjected to a second round of centrifugation under the same condition for 45 minutes. The cleared lysate was incubated with Anti-FLAG M2 agarose beads (Sigma-Aldrich) for 3 hours at 4□, using 100 μL beads per batch of cell culture. The suspension was loaded onto a Poly-Prep chromatography column (Bio-Rad) and washed with 150 mL wash buffer (50 mM HEPES pH 7.5, 75 mM NH_4_Cl, 20 mM MgCl_2_, 0.3 mM GDP•BeF_3_^-^). Bound ribosomal particles were incubated overnight with 2 mL elution buffer (50 mM HEPES pH 7.5, 75 mM NH_4_Cl, 20 mM MgCl_2_, 0.9 mM GDP•BeF_3_^-^, 2 mM DTT, 20 μg TEV protease) and eluted. The collected samples were pooled and concentrated using a spin concentrator (Vivaspin 500, 100,000 MWCO, Sartorius). The concentration of ribosomal particles was estimated by negative-staining microscopy. Fresh samples were used for cryo-EM grid preparation without prior freezing. For the BipA dataset, cells were cultured at 20□ for ribosomal particles purification.

### Cryo-EM grid preparation and data acquisition

Grids were prepared using a Vitrobot Mark IV (FEI) operated at 100% humidity and 8□. Prior to sample vitrification, holey carbon grids (Quantifoil R 2/2) were coated with a thin layer of freshly prepared carbon film using an ACE600 sputter coater (Leica) and glow-discharged for 30 s at medium RF power setting with a plasma cleaner (PDC-32G-2, Harrick Plasma). Of the eluted ribosomal samples (apparent concentration ∼400 nM), 3.3 μL was applied to each discharged grid and incubated for 1 minute inside the Vitrobot chamber before blotting (blot time: 1s; blot force: -1) and plunge-freezing into liquid ethane.

Cryo-EM data were collected on Titan Krios transmission electron microscope (FEI) equipped with an energy filter (20 eV slit width) operating at 300 kV. For the YihA dataset, a total of 4,000 micrographs were acquired using SerialEM ^112^ on a K2 Summit detector (Gatan) at a pixel size of 1.052 Å (magnification of x130,000) with a total dose of 60^-^/Å^2^ fractionated over 32 frames and a defocus range of -0.8 to -1.8 μm. For the EngA, ObgE and BipA datasets, micrographs were collected using EPU (Thermo Scientific) on a K3 Summit detector (Gatan) at a pixel size of 1.07 Å (magnification of x81,000) with a total dose of 60^-^/Å^2^ fractionated over 32 frames and a defocus range of -0.8 to -1.6 μm. Detailed parameters are listed in **Table S1**.

### Cryo-EM image processing

All cryo-EM data processing steps were performed using RELION 4.0 ^113^ unless otherwise specified, and are summarized in **Figures S2-S4 and S11**. For each dataset, images were motion corrected and dose weighted using MotionCor2 ^114^. The motion-corrected micrographs were used for contrast transfer function (CTF) estimation with GCTF ^115^. Particles were automatically picked by the Laplacian-of-Gaussian (LoG) filter auto-picking and the neural-network based picker Topaz ^116^. Reference-free 2D classification was carried out to sort useful particles from junk particles, which were then subjected to 3D refinement, followed by multiple rounds of 3D classification with global and local angular search. And UCSF-Chimera ^117^ was used to visualize and interpret the maps. 3D classes corresponding to junk or low-quality particles were discarded. Well-resolved classes were pooled and subjected to 3D refinement and CTF refinement (beam-tilt, per-particle defocus and per-micrograph astigmatism) in RELION 4.0, followed by Bayesian polishing. A second round of 3D refinement and CTF refinement (beam-tilt, trefoil, tetrafoil, magnification anisotropy, per-particle defocus and per-micrograph astigmatism) were performed, followed by 3D refinement to calculate the consensus maps.

To deal with the compositional and conformational heterogeneity present in the consensus maps, non-align focused 3D classifications were performed in RELION 4.0 using appropriate tau2_fudge and masks encompassing the factor or RNA segment of interest unless noted (**Figures S2-S4 and S11**). For the L1 stalk in the EngA dataset, focused 3D refinement with local angular search followed by three rounds of non-align focused 3D classifications was carried out on recentered and reboxed particles to resolve different conformational states (**Figure S15**). Particles corresponding to each well-defined state were subsequently subjected to 3D refinement using the appropriate solvent masks. To improve local resolution, focused 3D refinements with local angular search were performed using region-specific masks.

The nominal resolutions of all cryo-EM maps were estimated on the basis of the gold-standard Fourier shell correlation (FSC) at the 0.143 criterion implemented in CryoSPARC ^118^. Maps were corrected for the modulation transfer function of the detector, and subjected to B-factor sharpening with an empirically determined negative B-factor in RELION 4.0. Local resolution evaluations were determined in RELION 4.0 with two independently refined half-maps. All final cryo-EM maps were post-processed by EMready ^119^ for visualization and model building. Image processing statistics are summarized in **Table S1**.

### Model building, refinement and analysis

A combination of AlphaFold-predicted models ^120^, existing cryo-EM structures, and *de novo* model building was used to build the pre-50S assembly intermediates. The structures of pre-50S (PDB ID: 7BL3, 7BL5) ^121^ and mature 50S subunit (PDB ID: 6PJ6) ^122^ were used as initial templates, which were rigid-body docked into the maps using UCSF Chimera, morphed with distance, torsion and secondary structure restraints from high-resolution reference (PDB ID: 7K00), and manually adjusted in COOT ^123^. To build assembly factors, AlphaFold-predicted structures were used as starting models rigid-body fitted into the corresponding cryo-EM densities, morphed with self-restraints and further refined manually in COOT. For EngA^PTE^, the local region of the map showing the highest resolution features was used to build amino acid residues by trial and error. The resulting fragment was queried against the *E. coli* protein database using BLAST to identify candidate protein and facilitate main-chain tracing. Secondary structure fragments from the AlphaFold-predicted EngA model were subsequently used to complete model building of EngA^PTE^.

To build BipA-bound translating ribosome from the BipA dataset, the high-resolution structure of *E.coli* 70S ribosome (PDB ID: 7K00) ^58^ and the AlphaFold-predicted BipA model were docked into the cryo-EM map, morphed with self-restraints and manually refined in COOT. Due to limited local resolution, the nascent chain was built *de-novo* with poly-alanine.

For translating ribosome containing EngA^PTE^ from the EngA dataset, the high-resolution structure of *E. coli* 70S ribosome (PDB ID: 7K00) was rigid-body fitted into the consensus cryo-EM map, and morphed with self-restraints followed by manual adjustments in COOT. The built model of EngA^PTE^ was then docked into the map and merged with the ribosome part. For analysis of intermediate-resolution reconstructions of EngA^PTE^-bound translation states, each containing EF-Tu, EF-G and BipA, respectively, the structures of 70S-EF-Tu (PDB ID: 5AFI) ^124^, 70S-EF-G (PDB ID: 7SSD) ^125^ and 70S-BipA (this study) were rigid-body fitted into the corresponding cryo-EM maps using UCSF Chimera, followed by manual rigid-body adjustments in COOT.

All models were real-space refined using phenix.real_space_refine in PHENIX ^126^ with secondary structure, Ramachandran and side chain rotamer restraints. Final models were validated using MolProbity ^127^ inside of the PHENIX software suit. Model refinement statistics are summarized in **Table S1**. The maps and models were analyzed and visualized in COOT, Chimera and ChimeraX ^128^. Figures and movies were prepared using ChimeraX.

### Sucrose gradient fractionation and immunoblotting

The experiments were performed as previously described with modifications ^22^. Briefly, BipA-tagged BW25113 strain (::3xFLAG-TEV-StrepII-BipA::) was cultured at 20□ in 200 mL LB medium to an OD600 of 0.5-0.6. Chloramphenicol (250 μg/mL) was added 5 min before harvesting. Cells were collected by centrifugation, resuspended in lysis buffer (50 mM HEPES pH 7.5, 75 mM NH_4_Cl, 20 mM MgCl_2_, 250 μg/mL chloramphenicol, 25 μL RNase-free rDNase I, 25 μL RiboLock RNase inhibitor, 5 μg/mL lysozyme, 0.9 mM GDP•BeF_3_^-^, 1x protease inhibitor cocktail, 1 mM PMSF), and lysed by FastPrep homogenizer (MP Biomedicals). The resultant lysates were clarified by two rounds of centrifugation, and 15,000 A_260_ units of the cleared supernatant were loaded onto 10-40% (w/v) sucrose gradients prepared in the same buffer. Gradients were centrifuged at 39,000 rpm for 2.5 h at 4□ in a SW41-Ti rotor (Beckmann), followed by fractionation using the Biocomp Gradient Station (BioComp Instruments) with A_260_ as a readout. Fractions were subsequently analyzed by immunoblotting with anti-FLAG antibody (Sigma Aldrich). The A_260_ profiles were processed and visualized using Origin (OriginLab).

### Spotting assay and genetic complementation

For assessing the functional significance of the interaction between BipA and ribosomes, *E. coli* cells (wild-type, Δ*bipA*, and Δ*bipA* strains harboring pBAD plasmids expressing wild-type or mutant BipA) were cultured in LB medium at 37□ to the late exponential phase and subsequently diluted to an OD600 of 0.7. Six tenfold serial dilutions were prepared, and 5 μL of each dilution was spotted onto LB agar plates containing 0.1% (w/v) L-arabinose. The plates were incubated at 20□ until single colonies became visible. For assaying the genetic interaction between BipA and other noncanonical translation factors (EF-P, YeiP, UUP), Δ *bipA* strains were transformed with pBAD plasmids expressing wild-type BipA, EF-P, YeiP, UUP, or UUP mutant (PtIM or EQ2 variants). The same culture condition and plating procedure were followed as described above.

### Statistics and reproducibility

All attempts at reproducing the purification of native ribosomal particles were successful. Spotting assays were performed with at least three independent biological replicates (n=3) for each condition. Image processing was carried out using established algorithms and software packages with embedded statistical analyses. Resolution estimations of cryo-EM maps are based on the gold-standard Fourier Shell Correlation (FSC) of 0.143 criterion ^129,130^. No additional statistical analyses were applied in this work.

