## Supplementary material for "Bacterial GTPases act as successive placeholders to mediate ribosome assembly and its coupling to translation initiation": Supplemental Information.pdf


**This PDF file includes:**

Figures S1-S17

Table S1

Movies S1 and S2

### Supplemental Figure and Figure Legends

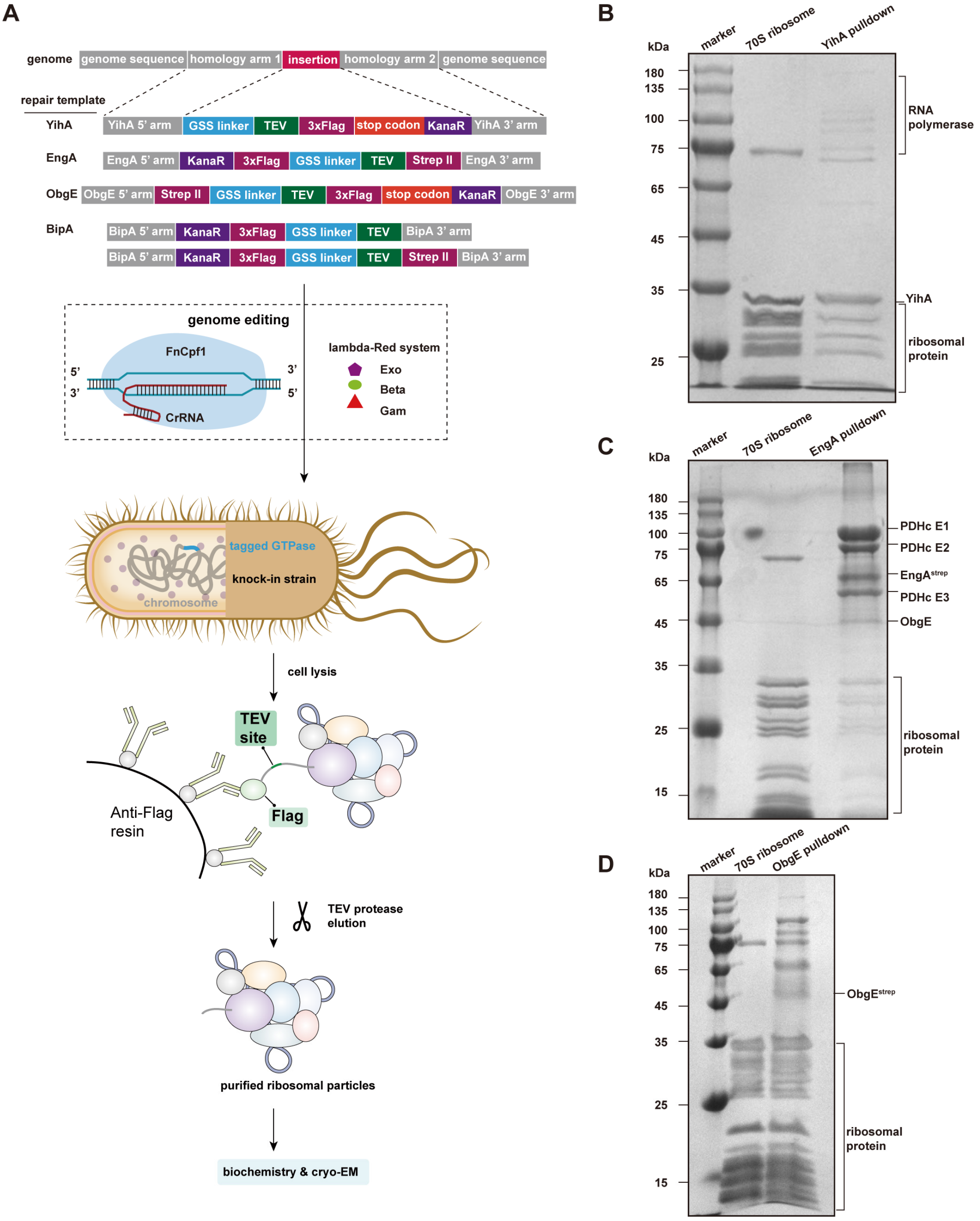


**Figure S1. Purification of endogenous bacterial ribosomal particles, related to Figure 1.**

(**A**) Strategy for the tagging of bacterial GTPases (YihA, EngA, ObgE, BipA) using the CRISPR and Lambda-Red system. Each repair template contains a pair of homology arms of the gene of interest, an affinity purification tag and a selectable resistance marker. One-step affinity purification was used to isolate native ribosomal particles.

(**B-D**) Coomassie blue-stained SDS-PAGE analysis of the purified samples through epitope tagged YihA (**B**), EngA (**C**), and ObgE (**D**).

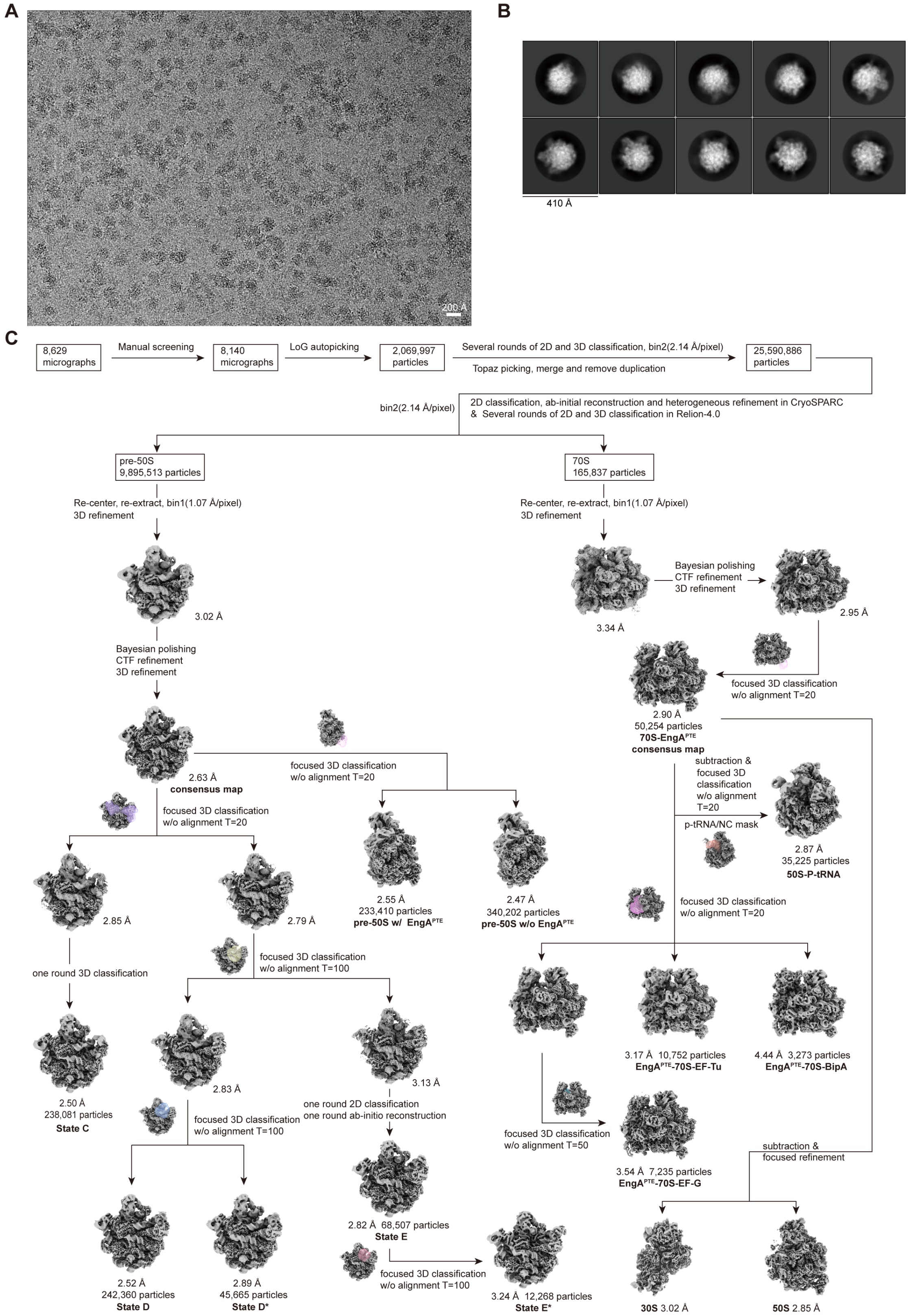


**Figure S3. Cryo-EM image processing of EngA dataset, related to Figures 1 and 6.**

(**A**) Motion-corrected representative cryo-EM micrograph.

(**B**) Representative 2D class averages.

(**C**) Data processing workflow.

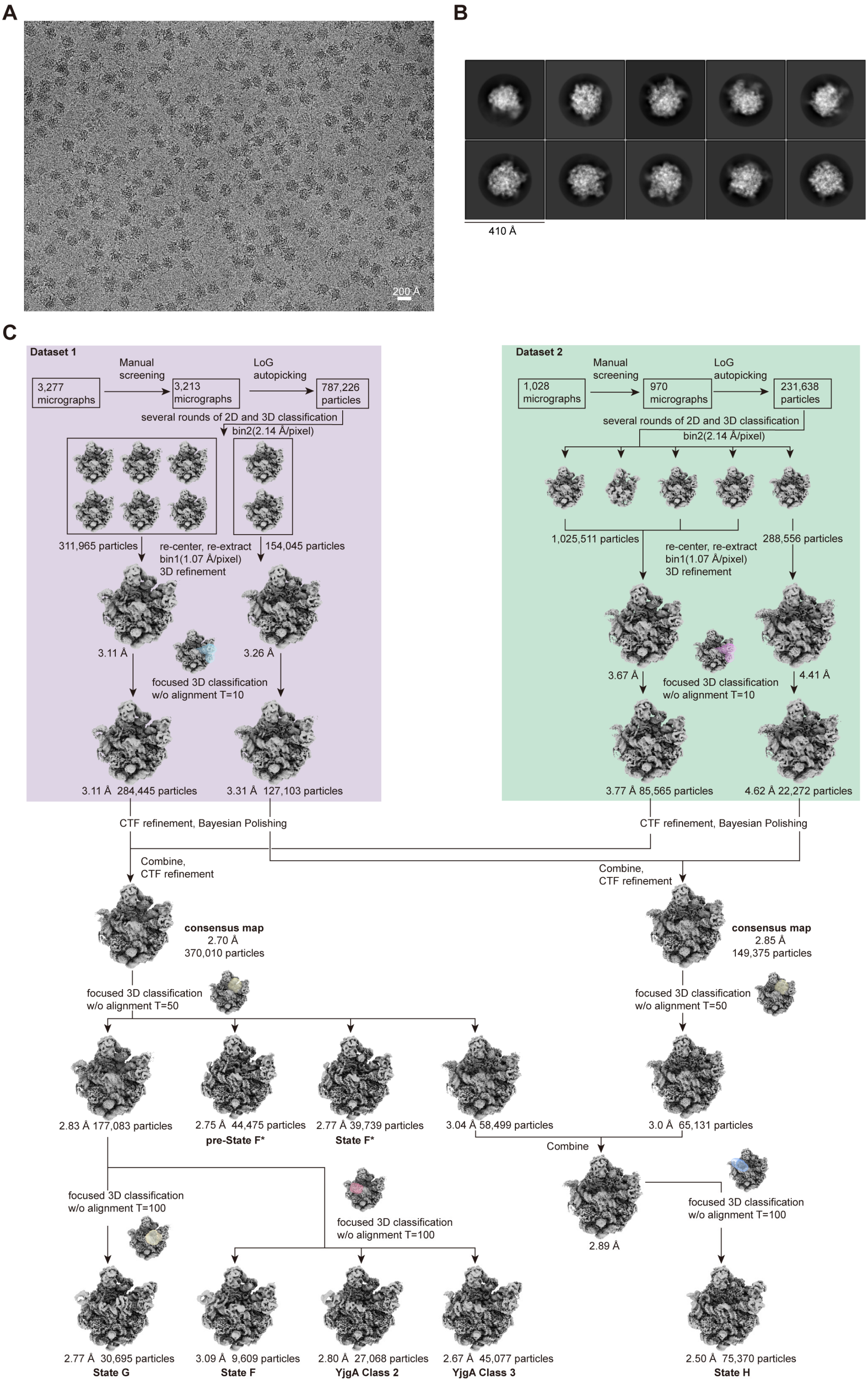


**Figure S4. Cryo-EM image processing of ObgE dataset, related to Figure 1.**

(**A**) Motion-corrected representative cryo-EM micrograph.

(**B**) Representative 2D class averages.

(**C**) Data processing workflow.

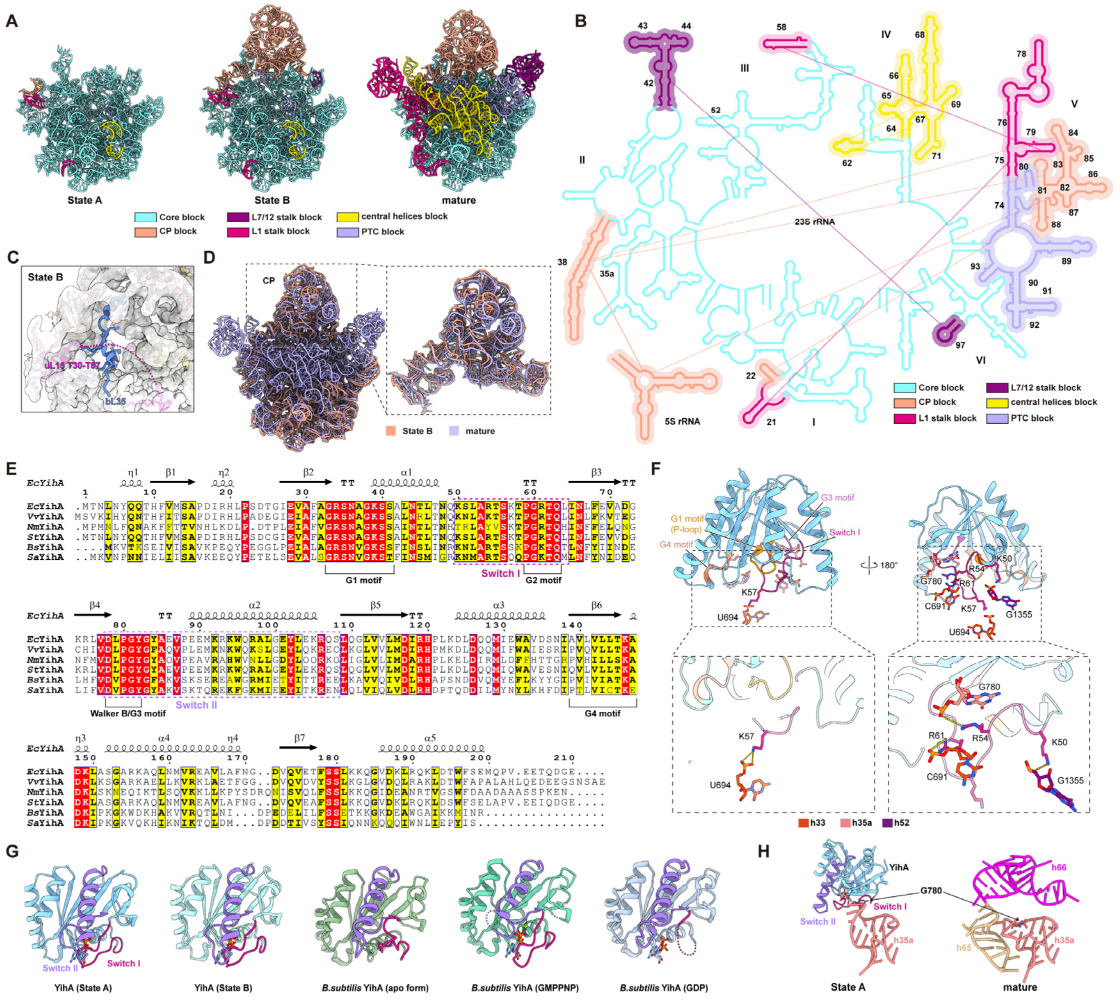


**Figure S5. Structural analysis of the YihA-mediated 23S rRNA maturation, related to Figure 2.**

(**A**) Comparison of the 23S rRNAs in States A, B and the mature 50S subunit (PDB ID: 6PJ6). RNA segments are colored according to the assigned blocks.

(**B**) Blocks of the 23S rRNA depicted in secondary structure diagram. Interactions among distal rRNA elements within each block are indicated by solid lines.

(**C**) Cryo-EM densities of the bL35 (blue) binding site in State B. Residues T30-T67 of uL15 (magenta) is disordered. bL35, from a previously published structure (PDB ID: 6PJ6), is superposed to show the absence of bL35 in State B.

(**D**) Comparison of the CP in State B and the mature 50S subunit (PDB ID: 6PJ6).

(**E**) Sequence alignment of YihA. *Ec, E. coli*; *Vv, V. vulnificus*; *Nm, N. meningitidis*; *St, S. typhimurium*; *Bs, B. subtilis*; *Sa, S. aureus*.

(**F**) Interactions of the Switch I (violet red) of YihA with rRNA helices h33, h35a and h52. Key residues are shown in stick representation.

(**G**) Cartoon representation of the structures of YihA (States A and B) and its *Bacillus subtilis* homolog (PDB ID: 1SUl, apo form; PDB ID: 1SVW, GMPPNP; PDB ID: 1SVI, GDP). The Switch I of YihA adopts a catalytically active conformation in our pre-50S complexes.

(**H**) Structural comparison of h35a in State A and the mature 50S subunit (PDB ID: 6PJ6).

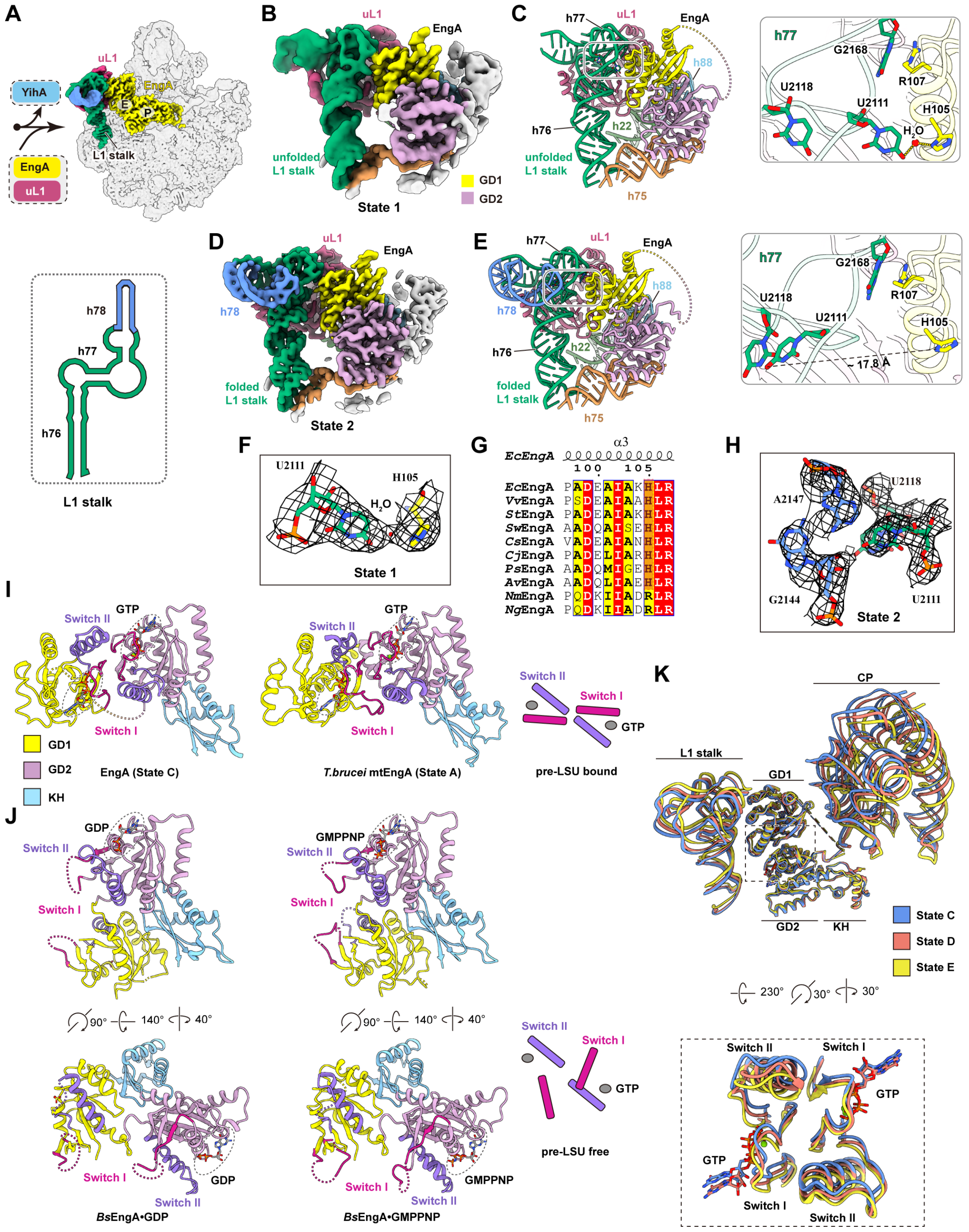


**Figures S6. EngA facilitates the maturation of the L1 stalk, related to Figure 3.**

(**A**) Cryo-EM map of State C, with the L1 stalk, EngA and uL1 color-coded. The L1 stalk acquires its rigidity first in State C.

(**B and C**) Cryo-EM map (**B**) and model (**C**) of the L1 stalk with disordered h78 in State 1.

(**D and E**) Cryo-EM map (**D**) and model (**E**) of the L1 stalk with fully folded h78 in

State 2. Key residues that show different conformations in State 1 and State 2 are shown in stick representation (right panels of **C** and **E**).

(**F**) Local density (contour level 6σ) of U2111 (h77) and H105 (EngA) in State 1. The potential water molecule is represented as a red sphere.

(**G**) Sequence alignment of the H105-containing helix of EngA in Gram-negative bacteria. H105 is highlighted in orange. *Ec*, *E. coli*; *Vv*, *V. vulnificus*; *St*, *S. Typhimurium*; *Sw*, *S. woodyi*; *Cs*, *C. salexigens*; *Cj*, *C. japonicus*; *Ps, P. syringae*; *Av, A. vinelandii*; *Nm, N. meningitidis*; *Ng, N. gonorrhoeae.*

(**H**) Local density (contour level 7σ) of U2111 (h77), U2118 (h77), G2144 (h78) and A2147 (h78) in State 2.

(**I and J**) Conformations of the Switch I and Switch II of EngA and its mitochondrial counterpart (PDB ID: 6YXX) when bound with pre-LSU (**I**), and its homolog in *Bacillus subtilis* (PDB ID: 5M7H, GMPPNP; 4DCS, GDP) free of pre-LSU binding (**J**). Domains of EngA are shown in cartoon representation: GD1 (yellow), GD2 (plum), KH (sky blue), Switch I (violet red) and Switch II (purple). Corresponding schematic diagrams depicting conformation of Switch I and Switch II are shown on the right panels.

(**K**) Conformational changes of EngA concomitant with the dynamics of the L1 stalk and CP. Structures of State C (blue), State D (tomato), and State E (yellow) are aligned by the KH domain of EngA. Zoom-in view for Switch I and Switch II is shown in the bottom panel.

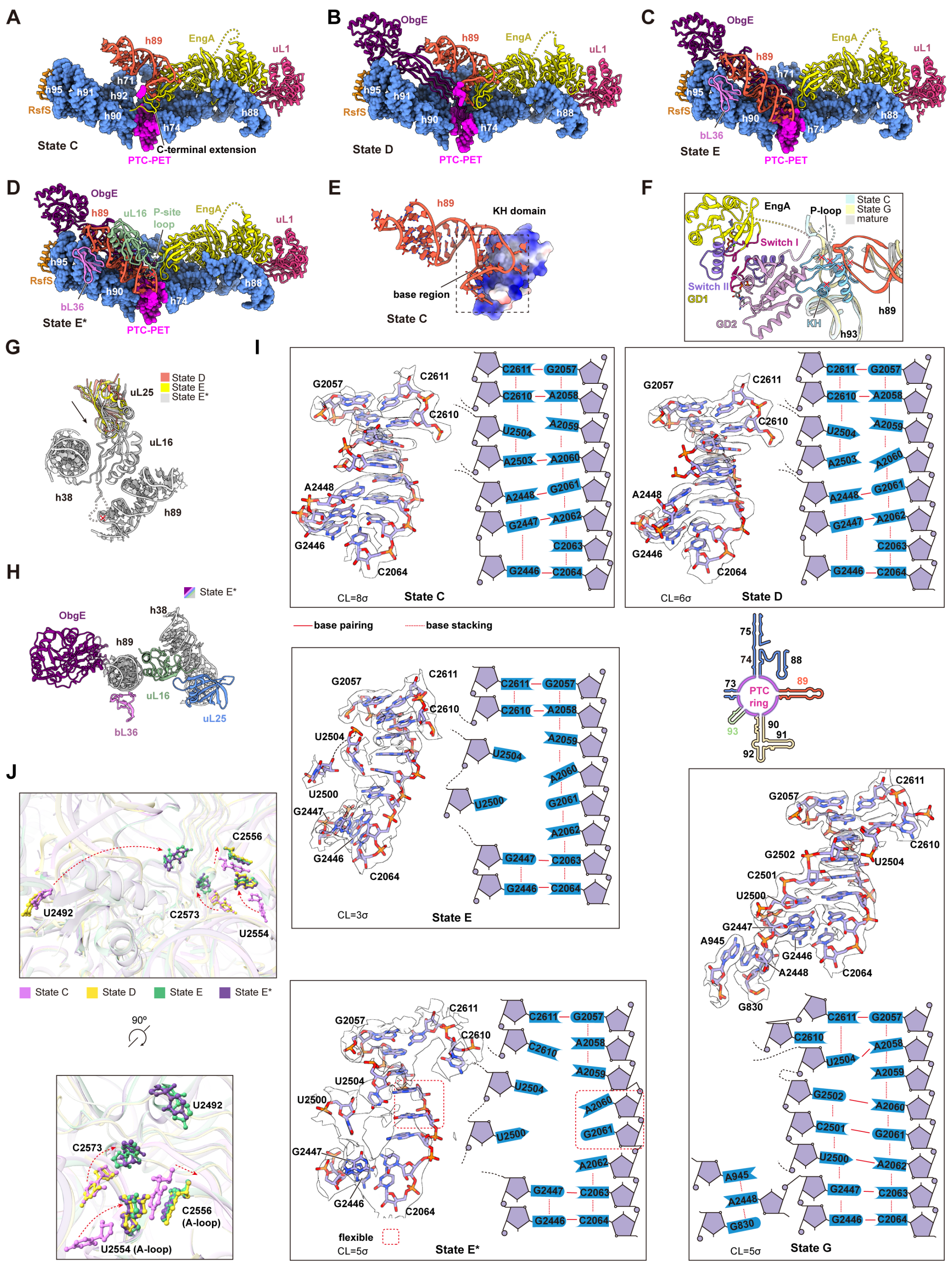

**Figure S7. Structural analyses of EngA- and ObgE-mediated maturation of the PTC region, related to Figure 4.**

(**A-D**) Structures of the PTC regions in States C (**A**), D (**B**), E (**C**), and E* (**D**). Components of the PTC are shown in solid surface representation (sky blue), h89 in cartoon representation (tomato), and the PTC-PET junction in solid surface representation (magenta). EngA, ObgE, uL1, RsfS, bL36, and uL16 are color-coded.

(**E**) Electrostatic surface representation of the KH domain of EngA in complex with h89 in premature conformation (tomato).

(**F**) Steric clash between the KH domain of EngA and the P-loop docking site in the mature 50S subunit. The immature P-loop (State G) and the mature one (PDB ID: 6PJ6) are superposed on State C for clarity. Clashed sites are indicated with red crosses.

(**G**) Continuous movements of uL25 facilitate the incorporation of uL16 in State E*.

(**H**) Structure of h89 in near mature conformation in State E*, clamped by bL36, uL16 and ObgE.

(**I**) Structural rearrangements of the PTC-PET junction from States C to E*. Corresponding elements in State G (near mature conformation) is also included for comparison. Nucleotides are shown in stick representation, with corresponding schematic diagrams provided for clarity.

(**J**) The maturation of the A-tRNA binding site. Nucleotides undergoing substantial conformational changes from States C to E* are depicted in stick representation.

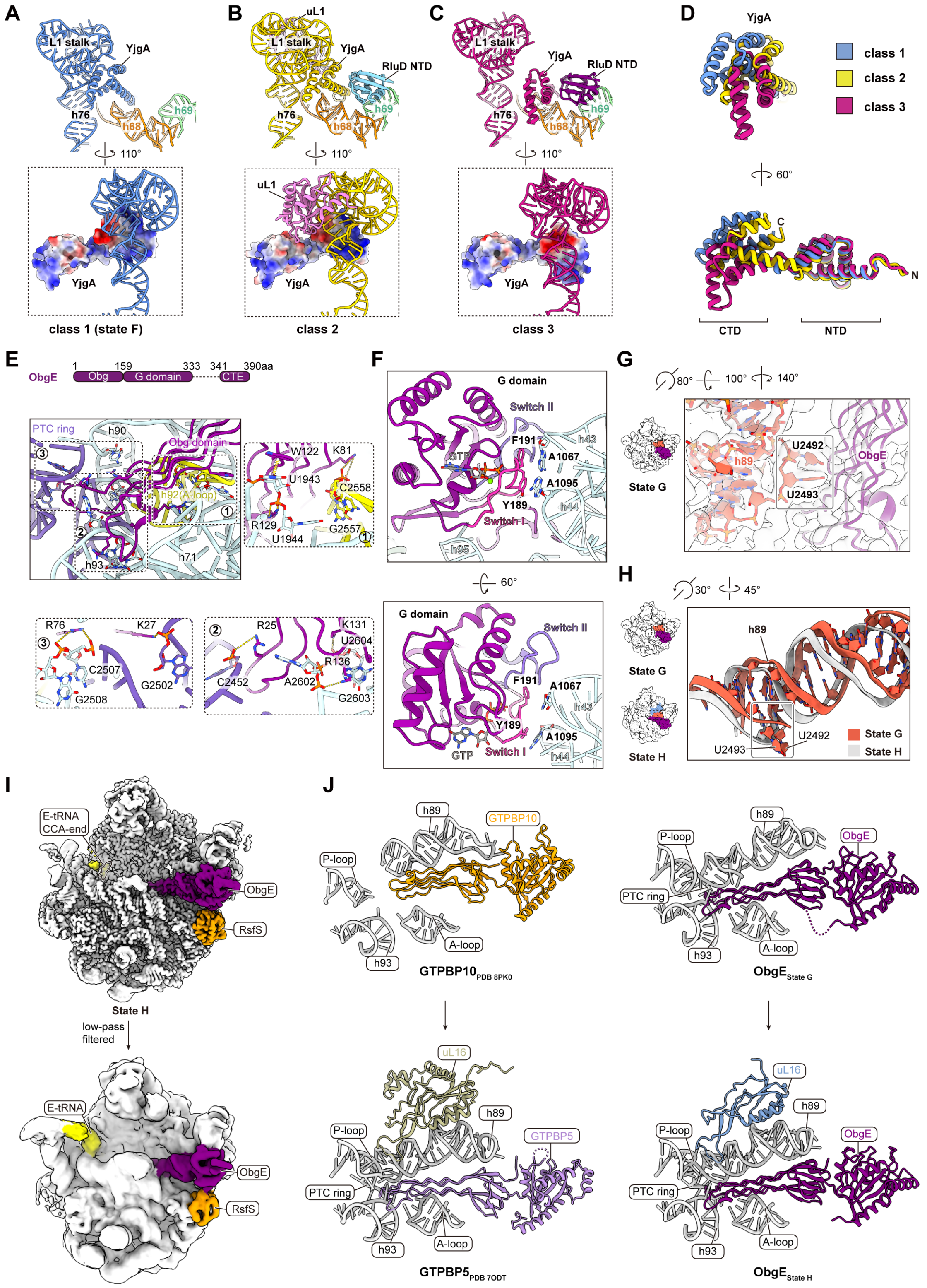

**Figure S8. Structural dynamics of YjgA and the interactions between ObgE and the 23S rRNA, related to Figure 5.**

(**A-C**) Three kinked conformational states of YjgA classified from the ObgE dataset: class 1, (**A**); class 2 (**B**); class 3 (**C**). YjgA is colored according to conformational states:

class 1 (blue); class 2 (yellow); class 3 (violet red). Electrostatic surface representation of YjgA is shown in the bottom panel.

(**D**) Structural comparison of YjgA from class 1-3.

(**E**) Interaction of the Obg domain of ObgE with the PTC region. Three close-up views (1-3) of the interactions are shown in the bottom and right panels. Key residues (K81, W122 and R129 in inset 1; R25, K131and R136 in inset 2; K27 and R76 in inset 3) of ObgE are shown in stick representation.

(**F**) Interaction of the G domain of ObgE with rRNA helices h43, h44 and h95. Switch I (pink) and Switch II (slate blue) are indicated.

(**G**) Close-up view of the interface between the immature h89 and the Obg domain in State G.

(**H**) Comparison of the premature h89 in State G with its mature state in State H.

(**I**) Cryo-EM map of State H. After low-pass filtration, the binding of the E-site tRNA becomes visible.

(**J**) Structural comparison of ObgE with its mitochondrial counterparts GTPBP10 (PDB ID: 8PK0) and GTPBP5 (PDB ID: 7ODT).

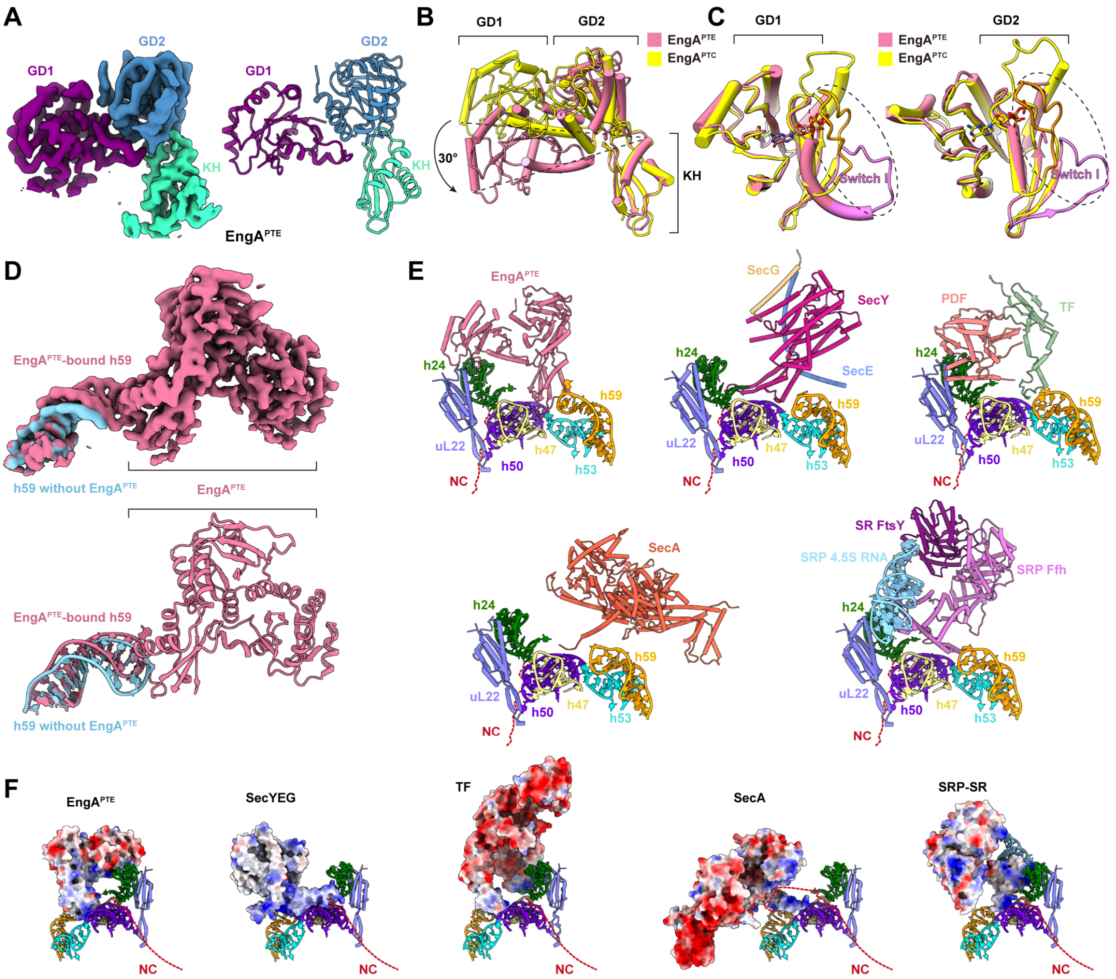


**Figure S9. Structure of EngA$^{PTE}$ and its comparison with other co-translation factors at the PTE, related to Figure 6.**

(**A**) Cryo-EM map and atomic model of EngA$^{PTE}$. GD1 (purple), GD2 (blue), KH (cyan).

(**B**) Structural comparison of EngA$^{PTE}$ (violet red) and EngA$^{PTC}$ (yellow).

(**C**) Comparison of GD1 and GD2 of EngA$^{PTE}$ (violet red) with that of EngA$^{PTC}$ (yellow).

(**D**) Comparison of the 23S rRNA h59 in the presence or free of EngA$^{PTE}$.

(**E and F**) Structural comparisons of EngA$^{PTE}$ with other co-translation factors in cartoon (**E**) and electrostatic surface (**F**) representations. SecYEG (PDB ID: 5GAE); PDF-TF (PDB ID: 7D6Z); SecA (PDB ID: 6S0K); SRP-SR (PDB ID: 5GAD). Red dash lines denote the assumed nascent peptide chain (NC). Factors and rRNA segments are individually color-coded.

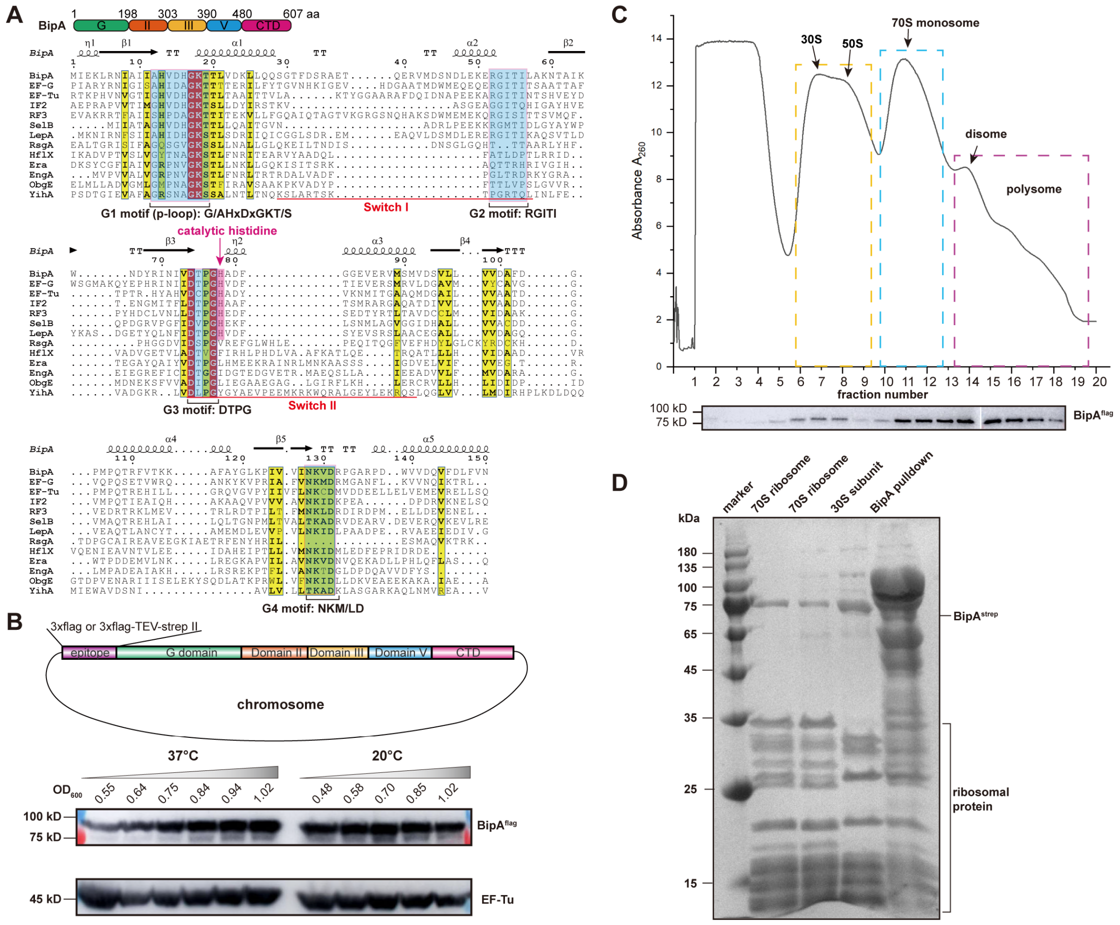


**Figure S10. Preparation and purification of endogenous ribosomal particles through epitope-tagged BipA, related to Figure 7.**

(**A**) Sequence alignment of the G domain from ribosome-associated GTPases (BipA, EF-G, EF-Tu, IF2, RF3, SelB, LepA, RsgA, HflX, Era, EngA, ObgE and YihA) in *E. coli*. Canonical catalytic histidine residue for GTP hydrolysis, conserved G domain motifs (G1-G4), and Switch I/II are indicated.

(**B**) Expression of endogenous BipA at normal (37 °C) or low (20 °C) growth temperature.

(**C**) Distribution of BipA across ribosomal sucrose gradient fractions at 20°C. Fraction peaks of the 30S subunit, 50S subunit, 70S ribosome and polysomes are indicated.

(**D**) Coomassie blue-stained SDS-PAGE analysis of the sample purified through epitope-tagged BipA.

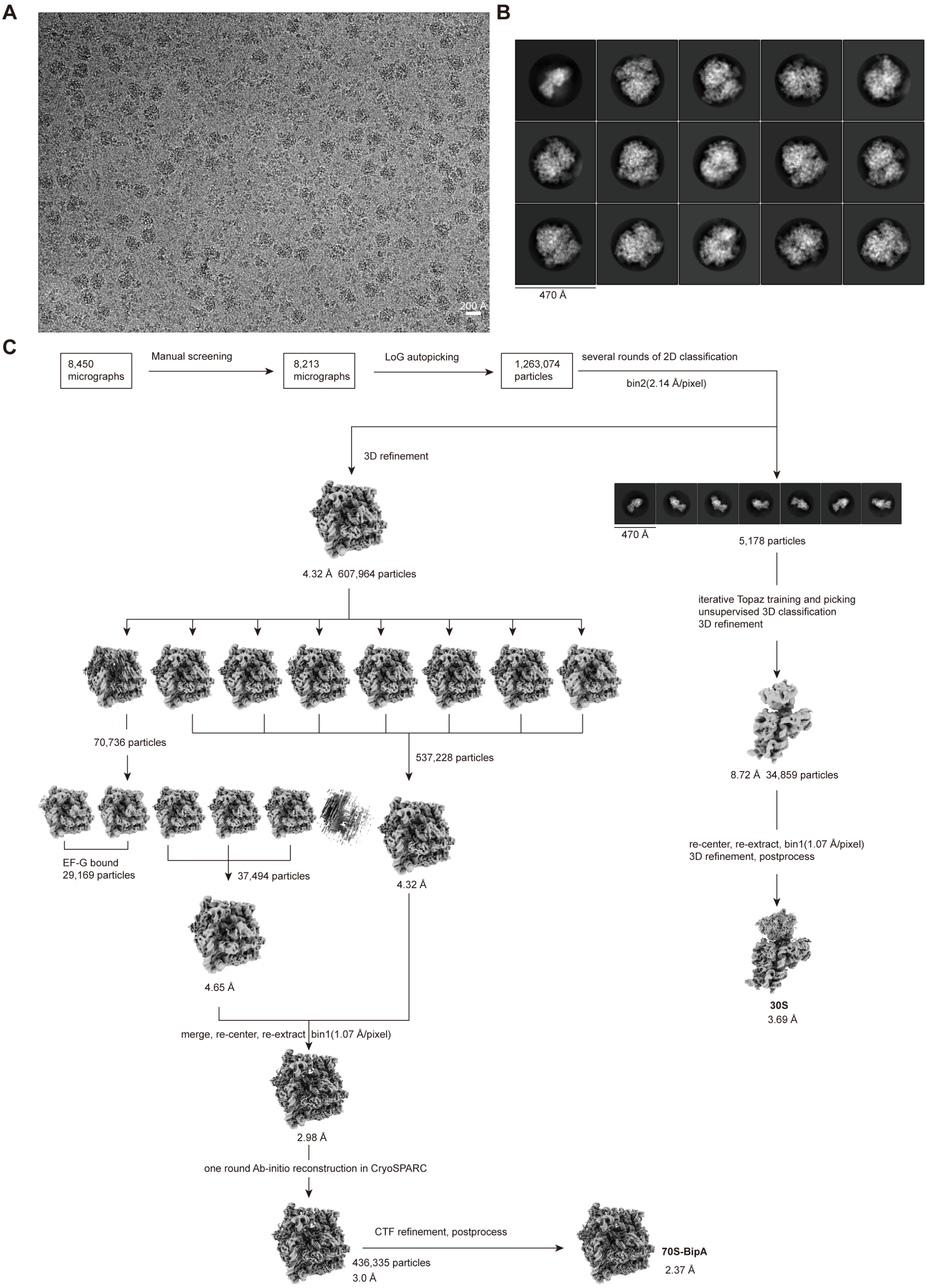


**Figure S11. Cryo-EM image processing of BipA dataset, related to Figure 7.**

(**A**) Motion-corrected representative cryo-EM micrograph.

(**B**) Representative 2D class averages.

(**C**) Data processing workflow.

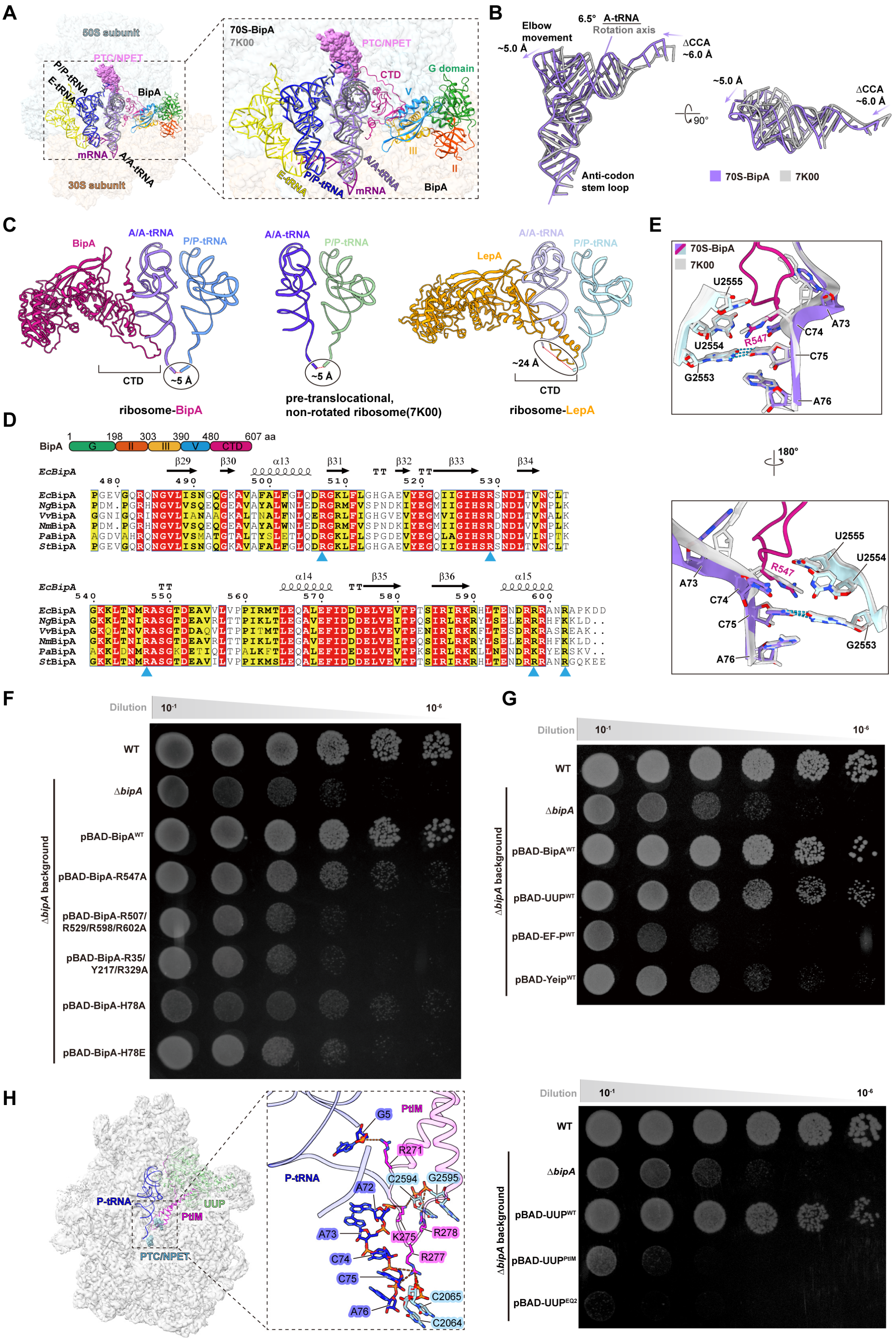


**Figure S12. Structural modulation of the A-site tRNA by BipA and functional analyses of the ribosome-BipA interfaces on cell growth, related to Figure 7.**

(**A**) Comparison of the A- and P-tRNAs in the 70S-BipA complex with that in canonical pre-translocational state (grey) (PDB ID: 7K00). The tRNAs are shown in zoom-in panel for clarity.

(**B**) Comparison of the A-site tRNA in the BipA-bound state with that in the canonical state (PDB ID: 7K00), aligned by anti-codon stem loop.

(**C**) Comparison of the geometry of the CCA-ends of the A- and P-tRNAs in different factor-bound states. Pre-translocational, non-rotated ribosome (PDB ID: 7K00); 70S-LepA (PDB ID: 3JCE).

(**D**) Sequence alignment of the CTD of BipA and its homologs among Gram-negative bacteria. Solid blue triangles indicate conserved residues interacting the A-site tRNA. *Ec*, *E. coli*; *Ng*, *N. gonorrhoeae*; *Vv*, *V. vulnificus*; *Nm*, *N. meningitidis*; *Pa*, *P. aeruginosa*; *St*, *S. Typhimurium*.

(**E**) Comparison of the CCA end of the A-site tRNA in the BipA-bound state with that in the canonical state (PDB ID: 7K00).

(**F**) Spot assay (growth at 20°C) analyses of BipA variants carrying mutations at the interfaces with the A-tRNA, the 30S subunit or at the active center of GTPase.

(**G**) Genetic complementation assay (growth at 20ºC) on EF-P, UUP and YeiP for rescuing the cold-sensitive phenotype of the Δ*bipA* strain.

(**H**) Spot assay (growth at 20ºC) analyses of the rescue of the Δ*bipA* phenotype by UUP. The P-tRNA structure (PDB ID: 7K00) and AlphaFold-predicted model of UUP were fitted into the cryo-EM map (EMDB ID: 29399) and manually adjusted for interaction analysis.

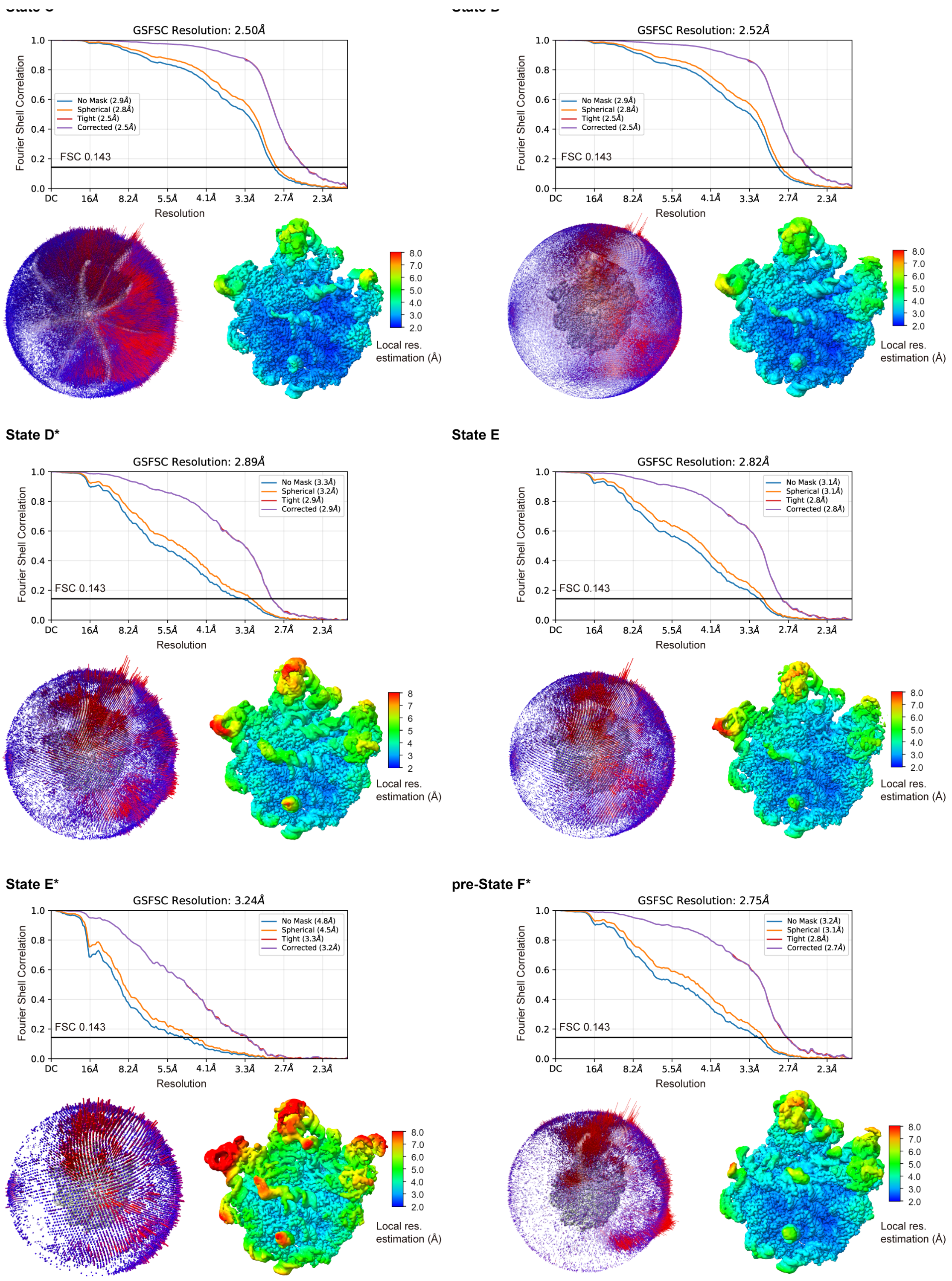


**Figure S13. Local resolution, orientation plots and FSC curves, related to Figure 1.**

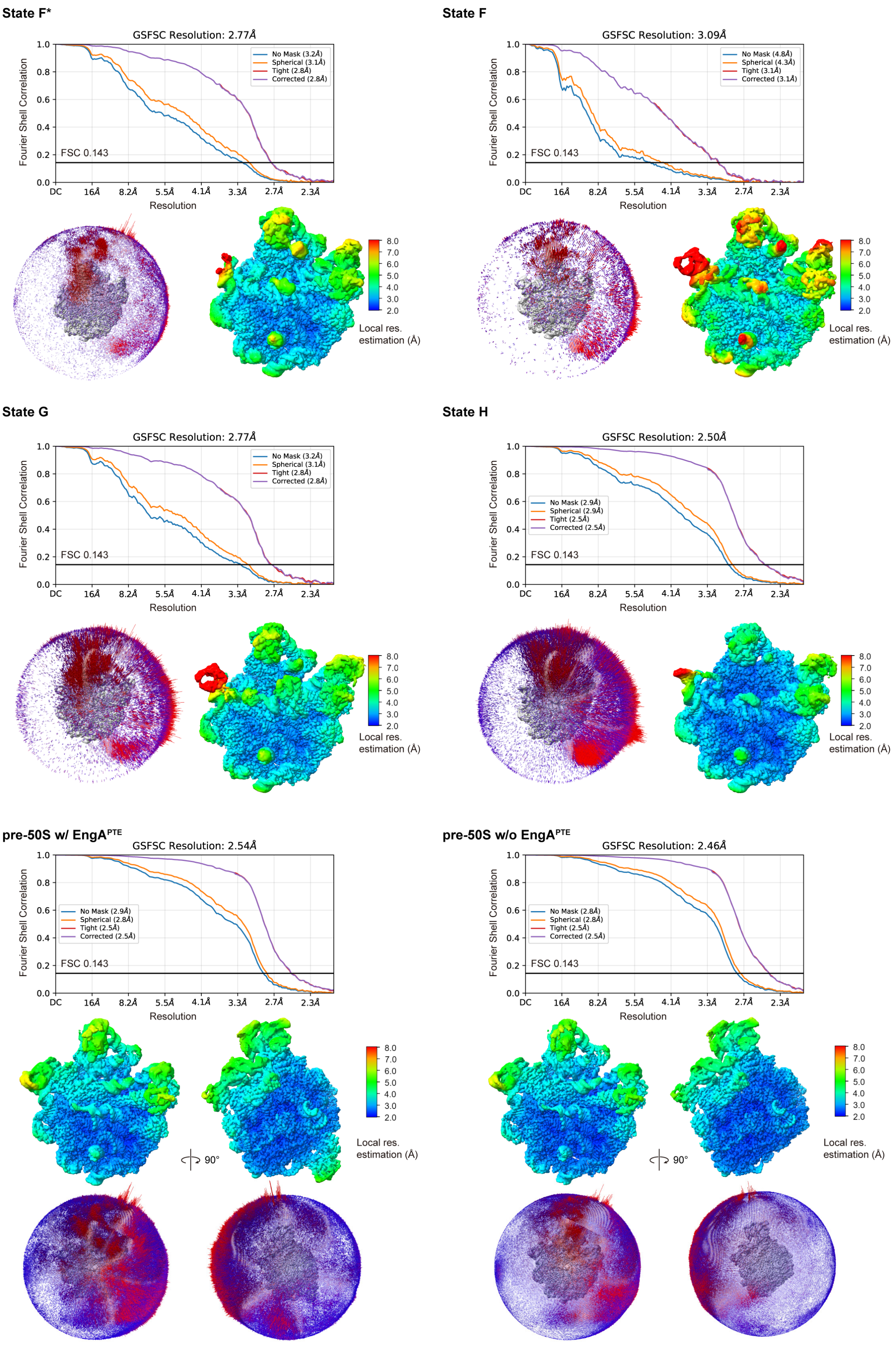


**Figure S14. Local resolution, orientation plots and FSC curves, related to Figures 1 and 6.**

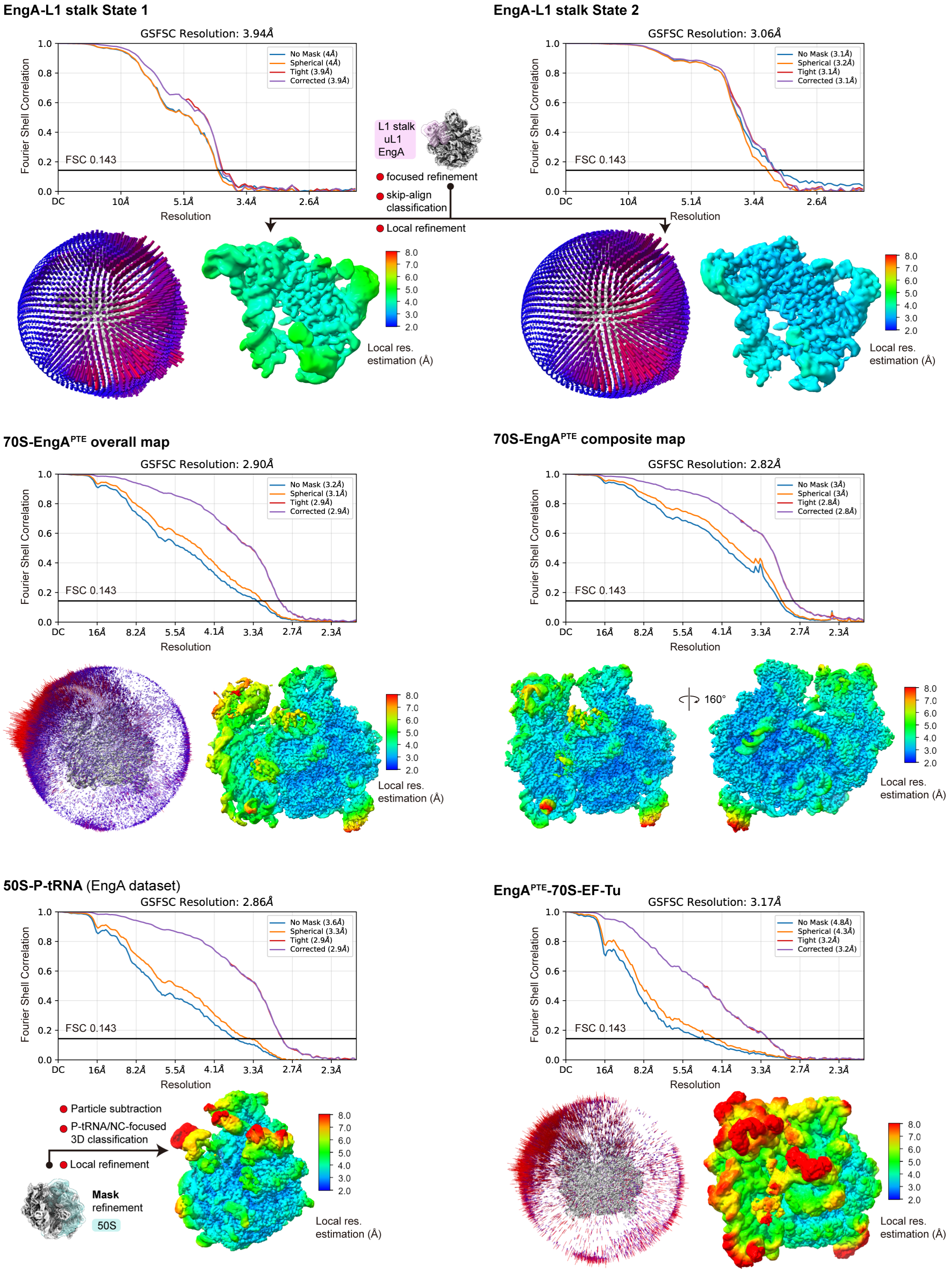


**Figure S15. Local resolution, orientation plots and FSC curves, related to Figures 1, 6 and S6.**

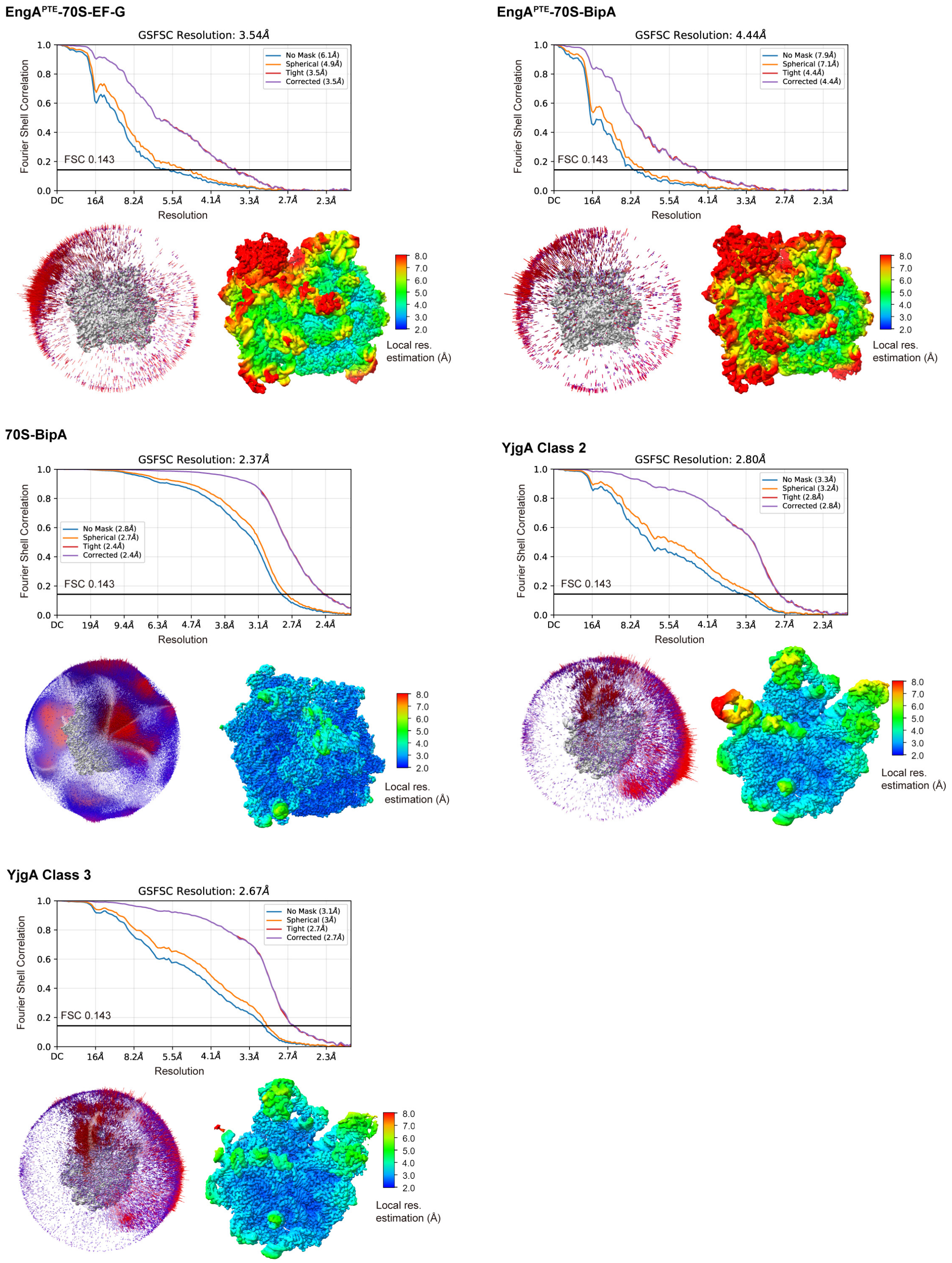


**Figure S16. Local resolution, orientation plots and FSC curves, related to Figures 6, 7 and S8.**

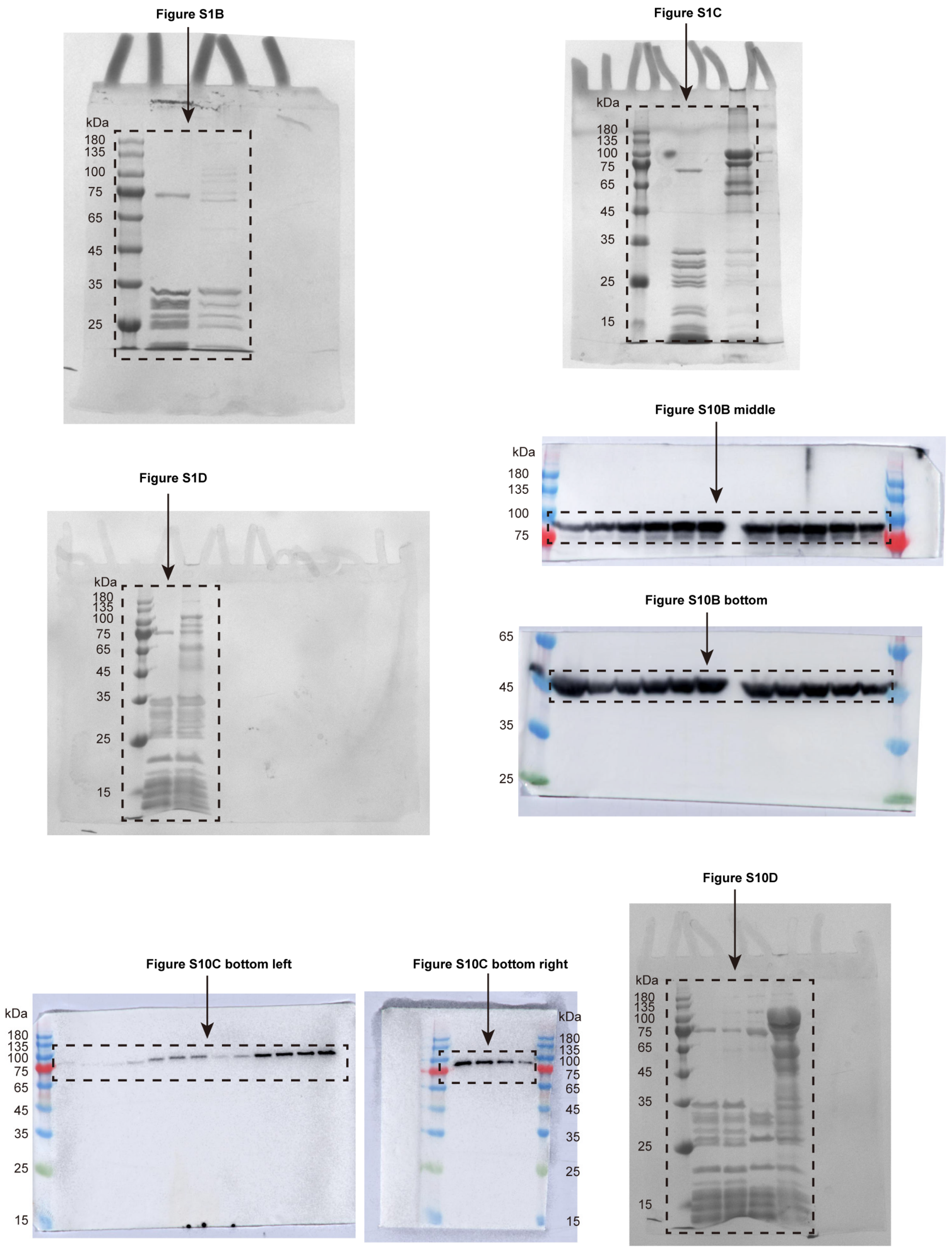


**Figure S17. Raw uncropped images of coomassie blue-stained gels and western blots, related to Figures S1 and S10.**

**Table S1. Cryo-EM data collection, refinement and validation statistics.**

| | State A (EMD-xxxxx) (PDB xxxx) | State B (EMD-xxxxx) (PDB xxxx) | State C (EMD-69028) (PDB 23JP) | State D (EMD-69031) (PDB 23JS) | State D* (EMD-69033) (PDB 23JU) | State E (EMD-69039) (PDB 23KB) |
|---|---|---|---|---|---|---|
| **Data collection and processing** | | | | | | |
| Microscope | Titan Krios | Titan Krios | Titan Krios | Titan Krios | Titan Krios | Titan Krios |
| Voltage (kV) | 300 | 300 | 300 | 300 | 300 | 300 |
| Camera | K2 Summit | K2 Summit | K3 Summit | K3 Summit | K3 Summit | K3 Summit |
| Magnification | 130,000 | 130,000 | 81,000 | 81,000 | 81,000 | 81,000 |
| Pixel size (Å/pixel) | 1.052 | 1.052 | 1.07 | 1.07 | 1.07 | 1.07 |
| Electron exposure ($e^-$/Å$^2$) | 60 | 60 | 60 | 60 | 60 | 60 |
| Number of frames | 32 | 32 | 32 | 32 | 32 | 32 |
| Defocus range (μm) | -0.8 to -1.8 | -0.8 to -1.8 | -0.8 to -1.6 | -0.8 to -1.6 | -0.8 to -1.6 | -0.8 to -1.6 |
| Energy filter slit width (V) | 20 | 20 | 20 | 20 | 20 | 20 |
| Micrographs collected (no.) | 4,000 | 4,000 | 8,629 | 8,629 | 8,629 | 8,629 |
| Initial particle images (no.) | | | 2,069,997 | 2,069,997 | 2,069,997 | 2,069,997 |
| Final particle images (no.) | | | 238,081 | 242,360 | 45,665 | 68,507 |
| Point-group symmetry | C1 | C1 | C1 | C1 | C1 | C1 |
| Map resolution (global, Å) | | | | | | |
| FSC 0.143 (masked) | | | 2.50 | 2.52 | 2.89 | 2.82 |
| Map resolution range (Å) | | | 2.19 – 6.66 | 2.19 – 6.4 | 2.57 – 8.21 | 2.46 – 8.16 |
| **Refinement** | | | | | | |
| Initial model used (PDB code) | 7K00 | 7K00 | 7BL3 | 7BL3 | 7BL3 | 7BL3 |
| Model resolution (Å) | 1.8 | 2.1 | 2.1 | 2.1 | 2.3 | 2.2 |
| FSC threshold | 0.5 | 0.5 | 0.5 | 0.5 | 0.5 | 0.5 |
| Model resolution range (Å) | | | | | | |
| Map sharpening *B* factor (Å$^2$) | | | -20.0 | -20.0 | -20.0 | -20.0 |
| Model composition | | | | | | |
| Non-hydrogen atoms | 53,240 | 66,070 | 89,853 | 93,308 | 93,626 | 95,014 |
| Protein residues | 1,818 | 2,335 | 3,679 | 4,125 | 4,138 | 4,307 |
| RNA | 1,814 | 2,223 | 2,864 | 2,869 | 2,878 | 2,884 |
| Ligands ($Mg^{2+}$/GTP) | 1/1 | 1/1 | 2/2 | 3/3 | 3/3 | 3/3 |
| Average *B* factors (Å$^2$) | | | | | | |
| Protein | 41.89 | 102.15 | 56.14 | 63.76 | 79.52 | 79.39 |
| RNA | 54.90 | 78.81 | 48.96 | 44.62 | 59.05 | 56.30 |
| Ligand | 56.12 | 88.72 | 46.22 | 64.98 | 79.99 | 95.05 |
| R.m.s. deviations | | | | | | |
| Bond lengths (Å) | 0.005 | 0.004 | 0.004 | 0.007 | 0.002 | 0.004 |
| Bond angles (°) | 0.656 | 0.591 | 0.589 | 0.710 | 0.541 | 0.603 |
| **Validation** | | | | | | |
| MolProbity score | 1.56 | 1.76 | 1.89 | 1.89 | 1.80 | 1.98 |
| Clashscore | 5.79 | 8.43 | 8.42 | 9.60 | 8.44 | 10.31 |
| Poor rotamers (%) | 0.67 | 0.00 | 1.38 | 0.09 | 0.15 | 0.20 |
| Ramachandran plot | | | | | | |
| Favored (%) | 96.40 | 95.62 | 95.27 | 94.37 | 95.03 | 92.92 |
| Allowed (%) | 3.55 | 4.38 | 4.59 | 5.58 | 4.89 | 7.01 |
| Disallowed (%) | 0.06 | 0.00 | 0.14 | 0.05 | 0.07 | 0.07 |

| | **State E*** (EMD-69048) (PDB 23KJ) | **pre-State F*** (EMD-69081) (PDB 23LT) | **State F*** (EMD-69084) (PDB 23LV) | **State F** (EMD-69087) (PDB 23MP) | **State G** (EMD-69101) (PDB 23NC) | **State H** (EMD-69110) (PDB 23NQ) |
|---|---|---|---|---|---|---|
| **Data collection and processing** | | | | | | |
| Microscope | Titan Krios | Titan Krios | Titan Krios | Titan Krios | Titan Krios | Titan Krios |
| Voltage (kV) | 300 | 300 | 300 | 300 | 300 | 300 |
| Camera | K3 Summit | K3 Summit | K3 Summit | K3 Summit | K3 Summit | K3 Summit |
| Magnification | 81,000 | 81,000 | 81,000 | 81,000 | 81,000 | 81,000 |
| Pixel size (Å/pixel) | 1.07 | 1.07 | 1.07 | 1.07 | 1.07 | 1.07 |
| Electron exposure ($e^-$/Å$^2$) | 60 | 60 | 60 | 60 | 60 | 60 |
| Number of frames | 32 | 32 | 32 | 32 | 32 | 32 |
| Defocus range (μm) | -0.8 to -1.6 | -0.8 to -1.6 | -0.8 to -1.6 | -0.8 to -1.6 | -0.8 to -1.6 | -0.8 to -1.6 |
| Energy filter slit width (V) | 20 | 20 | 20 | 20 | 20 | 20 |
| Micrographs collected (no.) | 8,629 | 4,305 | 4,305 | 4,305 | 4,305 | 4,305 |
| Initial particle images (no.) | 2,069,997 | 1,018,864 | 1,018,864 | 1,018,864 | 1,018,864 | 1,018,864 |
| Final particle images (no.) | 12,268 | 44,475 | 39,739 | 9,609 | 30,695 | 75,370 |
| Point-group symmetry | C1 | C1 | C1 | C1 | C1 | C1 |
| Map resolution (global, Å) | | | | | | |
| FSC 0.143 (masked) | 3.24 | 2.75 | 2.77 | 3.09 | 2.77 | 2.50 |
| Map resolution range (Å) | 2.92 – 10.92 | 2.37 – 7.60 | 2.41 – 7.28 | 2.85 – 10.84 | 2.42 – 9.98 | 2.21 – 9.80 |
| **Refinement** | | | | | | |
| Initial model used (PDB code) | 7BL3 | 7BL5 | 7BL5 | 7BL5 | 7BL5 | 6PJ6 |
| Model resolution (Å) | 2.5 | 2.3 | 2.2 | 2.4 | 2.1 | 2.0 |
| FSC threshold | 0.5 | 0.5 | 0.5 | 0.5 | 0.5 | 0.5 |
| Model resolution range (Å) | | | | | | |
| Map sharpening *B* factor (Å$^2$) | -20.0 | -15.0 | -15.0 | -15.0 | -15.0 | -15.0 |
| Model composition | | | | | | |
| Non-hydrogen atoms | 95,756 | 92,152 | 93,479 | 94,129 | 96,663 | 96,730 |
| Protein residues | 4,361 | 3,602 | 3,734 | 3,837 | 4,134 | 3,918 |
| RNA | 2,886 | 3,003 | 3,007 | 3,010 | 3,003 | 3,084 |
| Ligands ($Mg^{2+}$/GTP) | 1/3 | 0/1 | 1/1 | 0/1 | 1/1 | 130/1 |
| Average *B* factors (Å$^2$) | | | | | | |
| Protein | 113.63 | 65.34 | 68.08 | 85.28 | 71.53 | 48.06 |
| RNA | 82.70 | 63.42 | 65.55 | 64.33 | 68.76 | 54.33 |
| Ligand | 155.22 | 95.97 | 94.31 | 123.08 | 85.85 | 44.57 |
| R.m.s. deviations | | | | | | |
| Bond lengths (Å) | 0.003 | 0.004 | 0.003 | 0.003 | 0.004 | 0.006 |
| Bond angles (°) | 0.563 | 0.612 | 0.560 | 0.559 | 0.584 | 0.673 |
| **Validation** | | | | | | |
| MolProbity score | 1.97 | 1.73 | 1.70 | 1.82 | 1.80 | 1.82 |
| Clashscore | 10.60 | 8.27 | 7.67 | 9.43 | 8.47 | 7.91 |
| Poor rotamers (%) | 0.28 | 0.14 | 0.23 | 0.16 | 0.06 | 1.14 |
| Ramachandran plot | | | | | | |
| Favored (%) | 93.49 | 95.96 | 95.97 | 95.33 | 95.12 | 95.01 |
| Allowed (%) | 6.46 | 4.01 | 3.95 | 4.65 | 4.75 | 4.91 |
| Disallowed (%) | 0.05 | 0.03 | 0.08 | 0.03 | 0.12 | 0.08 |

| | **pre-50S w/ EngA$^{PTE}$** (EMD-69056) (PDB 23KW) | **pre-50S w/o EngA$^{PTE}$** (EMD-69053) (PDB 23KP) | **70S-BipA** (EMD-69017) (PDB 23JI) | **YjgA Class 2** (EMD-69117) (PDB 23NW) | **YjgA Class 3** (EMD-69120) (PDB 23NZ) | **70S-EngA$^{PTE}$** (EMD-69021) (PDB 23JJ) |
|---|---|---|---|---|---|---|
| **Data collection and processing** | | | | | | |
| Microscope | Titan Krios | Titan Krios | Titan Krios | Titan Krios | Titan Krios | Titan Krios |
| Voltage (kV) | 300 | 300 | 300 | 300 | 300 | 300 |
| Camera | K3 Summit | K3 Summit | K3 Summit | K3 Summit | K3 Summit | K3 Summit |
| Magnification | 81,000 | 81,000 | 81,000 | 81,000 | 81,000 | 81,000 |
| Pixel size (Å/pixel) | 1.07 | 1.07 | 1.07 | 1.07 | 1.07 | 1.07 |
| Electron exposure ($e^{-}$/Å$^2$) | 60 | 60 | 60 | 60 | 60 | 60 |
| Number of frames | 32 | 32 | 32 | 32 | 32 | 32 |
| Defocus range (µm) | -0.8 to -1.6 | -0.8 to -1.6 | -0.8 to -1.6 | -0.8 to -1.6 | -0.8 to -1.6 | -0.8 to -1.6 |
| Energy filter slit width (V) | 20 | 20 | 20 | 20 | 20 | 20 |
| Micrographs collected (no.) | 8,629 | 8,629 | 8,450 | 4,305 | 4,305 | 8,629 |
| Initial particle images (no.) | 2,069,997 | 2,069,997 | 1,263,074 | 1,018,864 | 1,018,864 | 2,069,997 |
| Final particle images (no.) | 233,410 | 340,202 | 436,335 | 27,068 | 45,077 | 50,254 |
| Point-group symmetry | C1 | C1 | C1 | C1 | C1 | C1 |
| Map resolution (global, Å) | | | | | | |
| FSC 0.143 (masked) | 2.54 | 2.46 | 2.37 | 2.80 | 2.67 | 2.90 |
| Map resolution range (Å) | 2.20 – 6.69 | 2.18 – 6.44 | 2.14 – 6.05 | 2.44 – 9.81 | 2.33 – 8.74 | 2.39 – 8.55 |
| **Refinement** | | | | | | |
| Initial model used (PDB code) | 7BL3 | 7BL3 | 7k00 | 7BL5 | 7BL5 | 7K00 |
| Model resolution (Å) | 2.1 | 2.1 | 1.6 | 2.2 | 2.2 | 2.5 |
| FSC threshold | 0.5 | 0.5 | 0.5 | 0.5 | 0.5 | 0.5 |
| Model resolution range (Å) | | | | | | |
| Map sharpening *B* factor (Å$^2$) | -20.0 | -25.0 | -25.0 | -15.0 | -15.0 | -25.0 |
| Model composition | | | | | | |
| Non-hydrogen atoms | 95,430 | 92,295 | 154,384 | 96,082 | 94,676 | 148,644 |
| Protein residues | 4,354 | 3,962 | 6,603 | 4,104 | 3,913 | 6,058 |
| RNA | 2,866 | 2,865 | 4,793 | 3,005 | 3,008 | 4,716 |
| Ligands ($Mg^{2+}$/GTP) | 4/4 | 3/3 | 1/1 | 1/1 | 1/1 | 0/1 |
| Average *B* factors (Å$^2$) | | | | | | |
| Protein | 65.57 | 62.27 | 37.54 | 70.91 | 58.66 | 119.46 |
| RNA | 47.47 | 49.35 | 37.44 | 43.46 | 66.54 | 94.78 |
| Ligand | 74.60 | 78.40 | 50.20 | 95.11 | 81.63 | 97.34 |
| R.m.s. deviations | | | | | | |
| Bond lengths (Å) | 0.004 | 0.002 | 0.003 | 0.005 | 0.003 | 0.004 |
| Bond angles (°) | 0.568 | 0.529 | 0.571 | 0.628 | 0.571 | 0.592 |
| **Validation** | | | | | | |
| MolProbity score | 1.76 | 1.68 | 1.31 | 1.75 | 1.68 | 1.81 |
| Clashscore | 8.06 | 7.31 | 4.22 | 8.15 | 7.47 | 7.99 |
| Poor rotamers (%) | 0.06 | 1.12 | 0.98 | 0.15 | 0.10 | 0.06 |
| Ramachandran plot | | | | | | |
| Favored (%) | 95.38 | 96.33 | 97.45 | 95.62 | 96.04 | 94.58 |
| Allowed (%) | 4.51 | 3.62 | 2.51 | 4.35 | 3.88 | 5.35 |
| Disallowed (%) | 0.12 | 0.05 | 0.03 | 0.03 | 0.08 | 0.07 |

| | **EngA$^{PTE}$-70S-BipA** (EMD-69018) | **EngA$^{PTE}$-70S-EF-G** (EMD-69019) | **EngA$^{PTE}$-70S-EF-Tu** (EMD-69020) | **EngA-L1 stalk State 1** (EMD-69140) (PDB 23PD) | **EngA-L1 stalk State 2** (EMD-69139) (PDB 23PC) | **50S-P-tRNA** (EngA Dataset) (EMD-69024) |
|---|---|---|---|---|---|---|
| **Data collection and processing** | | | | | | |
| Microscope | Titan Krios | Titan Krios | Titan Krios | Titan Krios | Titan Krios | Titan Krios |
| Voltage (kV) | 300 | 300 | 300 | 300 | 300 | 300 |
| Camera | K3 Summit | K3 Summit | K3 Summit | K3 Summit | K3 Summit | K3 Summit |
| Magnification | 81,000 | 81,000 | 81,000 | 81,000 | 81,000 | 81,000 |
| Pixel size (Å/pixel) | 1.07 | 1.07 | 1.07 | 1.07 | 1.07 | 1.07 |
| Electron exposure ($e^{-}$/Å$^2$) | 60 | 60 | 60 | 60 | 60 | 60 |
| Number of frames | 32 | 32 | 32 | 32 | 32 | 32 |
| Defocus range (μm) | -0.8 to -1.6 | -0.8 to -1.6 | -0.8 to -1.6 | -0.8 to -1.6 | -0.8 to -1.6 | -0.8 to -1.6 |
| Energy filter slit width (V) | 20 | 20 | 20 | 20 | 20 | 20 |
| Micrographs collected (no.) | 8,629 | 8,629 | 8,629 | 8,629 | 8,629 | 8,629 |
| Initial particle images (no.) | 2,069,997 | 2,069,997 | 2,069,997 | 2,069,997 | 2,069,997 | 2,069,997 |
| Final particle images (no.) | 3,273 | 7,235 | 10,752 | 77,910 | 656,889 | 35,225 |
| Point-group symmetry | C1 | C1 | C1 | C1 | C1 | C1 |
| Map resolution (global, Å) | | | | | | |
| FSC 0.143 (masked) | 4.44 | 3.54 | 3.17 | 3.94 | 3.06 | 2.85 |
| Map resolution range (Å) | 3.19 – 12.48 | 2.94 – 12.04 | 2.85 – 12.42 | 3.68 – 5.24 | 3.0 – 4.06 | 2.59 – 11.66 |
| **Refinement** | | | | | | |
| Initial model used (PDB code) | Not modeled (70S-BipA rigid-body fitted) | Not modeled (7SSD rigid-body fitted) | Not modeled (5AFI rigid-body fitted) | 7BL3 | 7BL3 | Not modeled (3JBU rigid-body fitted) |
| Model resolution (Å) | | | | 3.3 | 2.2 | |
| FSC threshold | | | | 0.5 | 0.5 | |
| Model resolution range (Å) | | | | | | |
| Map sharpening *B* factor (Å$^2$) | | | | -175.0 | -75.0 | |
| Model composition | | | | | | |
| Non-hydrogen atoms | | | | 7,786 | 8,339 | |
| Protein residues | | | | 580 | 579 | |
| RNA | | | | 155 | 181 | |
| Ligands ($Mg^{2+}$/GTP) | | | | | | |
| Average *B* factors (Å$^2$) | | | | | | |
| Protein | | | | 112.85 | 84.25 | |
| RNA | | | | 199.12 | 112.77 | |
| Ligand | | | | 89.07 | 83.28 | |
| R.m.s. deviations | | | | | | |
| Bond lengths (Å) | | | | 0.002 | 0.004 | |
| Bond angles (°) | | | | 0.523 | 0.544 | |
| **Validation** | | | | | | |
| MolProbity score | | | | 1.86 | 1.84 | |
| Clashscore | | | | 12.14 | 8.43 | |
| Poor rotamers (%) | | | | 0.21 | 0.00 | |
| Ramachandran plot | | | | | | |
| Favored (%) | | | | 96.17 | 94.42 | |
| Allowed (%) | | | | 3.83 | 5.58 | |
| Disallowed (%) | | | | 0.00 | 0.00 | |

**Movie S1. Coordinated movement of h89, h91, h95 and ObgE during progression from State G to State H, related to Figure 5.**

Progression from State G to State H involves coordinated movement of helices h89, h91 and h95, together with the Obg domain of ObgE, which would relieve the steric hindrance imposed on h89 by ObgE and induce conformational changes in Switch I and Switch II in the G domain, thus activating GTPase activity and promoting dissociation of ObgE from the pre-50S particles. The movie was prepared by morphing from State G to State H (gray).

**Movie S2. Density of SD-anti-SD duplex in the 70S-BipA complex, related to Figure 7.**

At the mRNA exit channel in the 70S-BipA complex, the density of SD-anti-SD duplex is observed, and is clamped by ribosomal protein bS21.
